# The CHIP–LMO7–BAG5 complex controls tau clearance and yields repurposed and newly designed therapeutic candidates for Alzheimer’s disease

**DOI:** 10.64898/2026.06.26.734369

**Authors:** Jiyoen Kim, Bakhos Tadros, Bhupesh Vaidya, Yan Hong Liang, Sung Yun Jung, Xue Deng, Jin Xu, Huda Yahya Zoghbi

## Abstract

Through cross-species genetic screens, we identified LMO7 as a regulator of tau levels *in vivo*. LMO7 bridges the E3 ligase CHIP and its inhibitory cochaperone BAG5, suppressing CHIP’s ligase activity and limiting tau ubiquitination and degradation. Adult LMO7 knockdown enhances CHIP activity, reduces total and phosphorylated tau, attenuates gliosis, and rescues memory deficits in tauopathy mice. Guided by AlphaFold-derived structural models and AI-based virtual screening, we developed a CHIP-derived competing peptide and identified FDA-approved drugs that disrupt the complex. Viral delivery of the peptide or oral administration of Telmisartan or MPA-2, an optimized Mycophenolic acid derivative, reduced pathogenic tau without toxicity. These findings establish CHIP–LMO7–BAG5 complex as a druggable node in tau homeostasis, with implications for tauopathies including Alzheimer’s disease, and *STUB1*-associated ataxias.

## Introduction

Tauopathies are a heterogeneous group of neurodegenerative disorders in which the accumulation of misfolded and aggregated tau is a major disease driver^1,2^. Alzheimer’s disease (AD), the most common tauopathy and the leading cause of dementia ranks among the most common causes of death among older adults^2^.The recent FDA approvals of anti-amyloid antibodies have validated disease-modification as a clinical goal but underscore the need to target tau, the proteinopathy that correlates most closely with cognitive decline and neuronal loss^3,4^. Multiple lines of evidence converge on tau-lowering as a tractable disease-modifying strategy. Genetic reduction of endogenous tau ameliorates amyloid-β–induced deficits in amyloid precursor protein (APP) transgenic mice^5^; a human carrying autosomal-dominant presenilin mutation and having amyloid β accumulation escapes cognitive decline in the absence of tau accumulation in the medial temporal lobe^6^; and our group has shown that even modest (∼20%) reductions in tau rescue pathology and cognition in tauopathy mice^7–10^. Translational momentum has accelerated: tau-targeted antisense oligonucleotides reduced soluble and aggregated tau in PS19 mice^11^ and lowered CSF tau in a Phase 1b trial of mild AD^12^ establishing tau-lowering as a clinically validated therapeutic axis^13^. A complementary, biologically distinct strategy — reactivating the brain’s own tau-clearance machinery — could broaden the therapeutic window and avoid the delivery and cost limitations of nucleic-acid drugs^14^.

Within the cellular proteostasis network, the U-box E3 ubiquitin ligase Carboxy-terminus of Hsp70-interacting protein (CHIP/STUB1) is a master regulator of misfolded-protein clearance^15,16^. CHIP polyubiquitinates phosphorylated and misfolded tau for proteasomal degradation^17–19^, and its substrate breadth extends across many proteins driving neurodegeneration: α-synuclein in Parkinson’s disease (PD)^20^, ataxin-1 in spinocerebellar ataxia type 1 (SCA1)^21,22^, and additional disease drivers^23–25^. Furthermore, loss-of-function variants in *STUB1* cause the recessive ataxia SCAR16^22,26,27^, and *STUB1* haploinsufficiency causes SCA48^28^, proving that even partial loss of CHIP activity is sufficient to drive human neurodegeneration. Pharmacologic activation of CHIP, however, has remained difficult: E3 ligases are notoriously hard to drug directly^29^, and known endogenous inhibitors of CHIP, notably Bcl-2 associated anthanogene 2 and 5 (BAG2 and BAG5), act through poorly understood mechanisms^20,30–32^.

Using cross-species genetic screens in human cells and *Drosophila*, our group previously identified several regulators of tau homeostasis, including the kinases NUAK1^8^ and TYK2^7^. Surprisingly, when we dissected the mechanisms of three additional, unbiasedly selected regulators (USP7, RNF130, or RNF149), all three converged on CHIP: USP7 deubiquitinates the CHIP-targeted lysines on tau, while RNF130 and RNF149 destabilize CHIP itself^10^. NUAK1, in turn, phosphorylates tau at Ser356 and disrupts CHIP recognition^8^. Together, these data position CHIP-mediated tau degradation as the pivotal node in tau homeostasis.

In the same screens, LIM domain only 7 (*LMO7*) emerged as a positive regulator of tau levels^10,33^. LMO7 has been implicated in actin binding and adherens-junction biology^34^ and in ubiquitin-pathway functions^35–37^, but no prior study has linked LMO7 to tau, to CHIP, or to neurodegeneration. Here show that LMO7 is the missing scaffold that permits BAG5 to inhibit CHIP. LMO7 binds CHIP and recruits BAG5 to form a CHIP–LMO7–BAG5 complex; disrupting this complex genetically or pharmacologically reactivates CHIP and lowers tau *in vivo*. Guided by AlphaFold-derived structural models, we designed a CHIP-derived competing peptide that disrupts the complex and lowers tau in mouse brain. We then used an AI-driven virtual screen of FDA-approved drugs^38^ and identified novel small-molecule disruptors that reduce pathogenic tau in PS19 mice without cytotoxicity. Our findings establish the CHIP–LMO7–BAG5 complex as a druggable, generalizable node for E3-ligase reactivation, with translational implications for AD, the STUB1 ataxias, and other CHIP-associated proteinopathies.

## Results

### LMO7 stabilizes tau by inhibiting CHIP E3 ubiquitin ligase activity

Having identified LMO7 was a positive regulator of tau^10^, we proceeded to validate the effect of LMO7 on tau levels in human cells and the mouse brain. We found that shRNA knockdown of LMO7 in human HEK293T cells reduced tau levels (**Fig. 1A-i**), while lentiviral overexpression of human LMO7 increased tau levels (**Fig. 1A-ii**). In mice, AAV-mediated *Lmo7* knockdown through intracerebroventricular injection at postnatal day zero (P0-ICV injection) also decreased tau levels in the brain (**Fig. 1B**). These findings indicate that LMO7 stabilizes tau in both human cells and the mouse brain.

**Figure 1:**
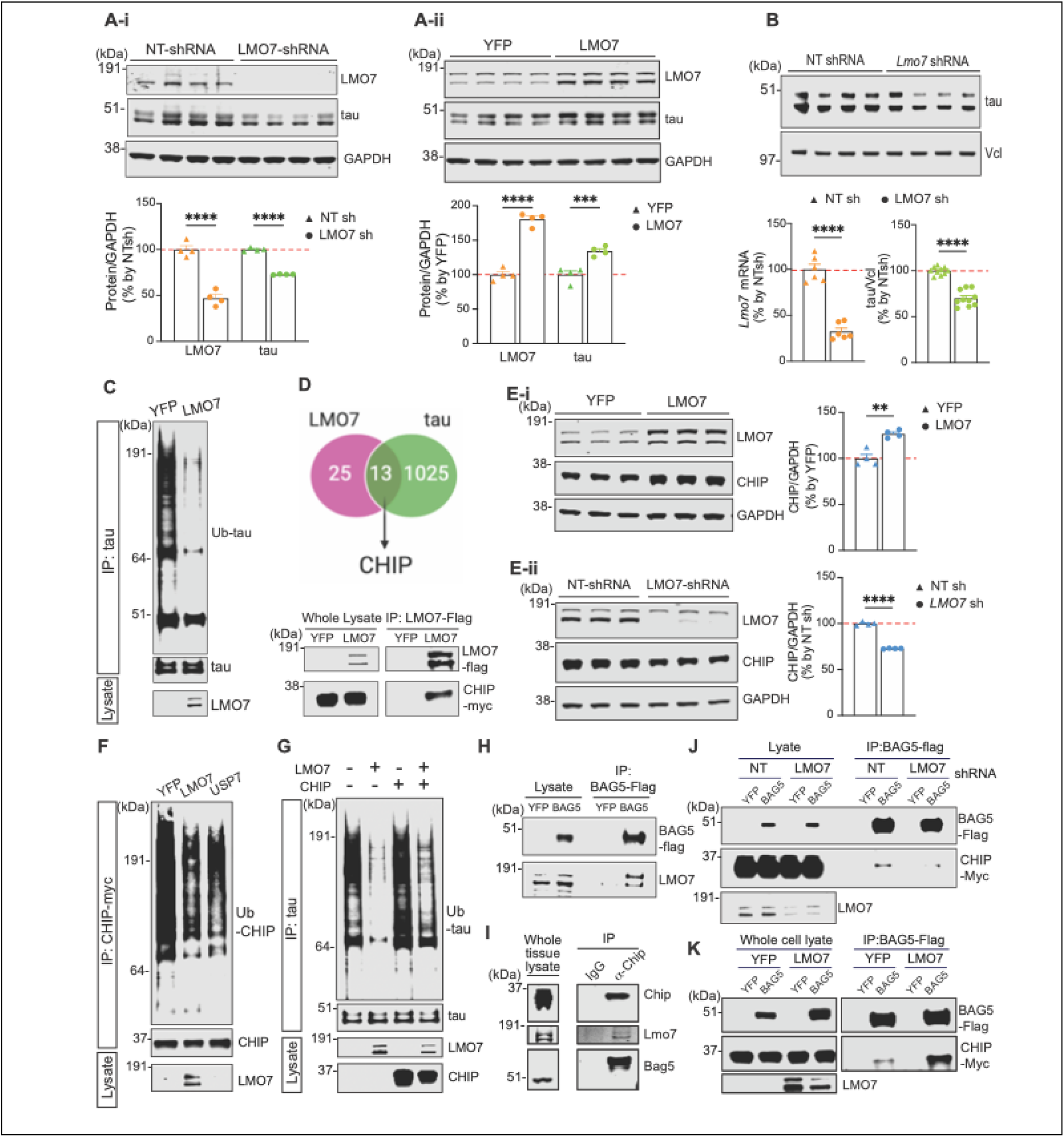
LMO7 antagonizes CHIP by mediating its interaction with the inhibitor, BAG5, thereby stabilizing tau. **(A)** Immunoblot (IB) image and quantification graph of tau and LMO7 level in HEK293T cells expressing either control non-targeting (NT) shRNA or *LMO7* shRNA **(A-i)**, or overexpressing YFP or LMO7 **(A-ii)**. **(B)** IB image and quantification graph of tau protein levels in the brains of wild-type mice injected with NT shRNA or *Lmo7* shRNA. Right panel: Quantification graph of *Lmo7* mRNA measured by qRT-PCR. **(C)** IB image of total and polyubiquitinated tau in HEK293T cell overexpressing either YFP or LMO7. **(D)** Venn diagram showing 13 shared interactors between LMO7 interactors (identified by our IP-MS) and tau interactors (listed in the BioGRID database), and IB image showing CHIP co-IP by LMO7 IP in HEK293T cells (lower panel). **(E)** IB image of CHIP and LMO7 in HEK293T cells expressing either YFP, LMO7 **(E-i)**, NT shRNA or *LMO7* shRNA **(E-ii)**. **(F)** IB image of total and ubiquitinated CHIP from cells expressing either YFP, LMO7, or the known regulators of CHIP-ubiquitination (USP7). **(G)** IB images of total- and ubiquitinated tau in the presence of either LMO7, CHIP, neither or both. **(H)** Co-IP of CHIP by BAG5-flag protein using anti-Flag antibody in HEK293T cells co-expressing CHIP-Myc with either YFP or BAG5-Flag. **(I)** IB image showing co-IP of Lmo7 and Bag5 by Chip IP in mouse brain tissue. **(J)** Co-IP of BAG5-Flag by CHIP-Myc with either NT shRNA or LMO7 shRNA in HEK293T cells. **(K)** Co-IP of BAG5-Flag by CHIP-Myc IP in HEK293T cells expressing either YFP or LMO7 proteins. Data shown as mean ± SEM (**p≤0.01, ***p≤0.001, ****p≤ 0.0001).

To investigate how LMO7 regulates tau, we next examined the effect of LMO7 on tau ubiquitination, given previous studies implicating LMO7 in ubiquitin signaling^35,36^. LMO7 overexpression markedly decreased the level of tau-polyubiquitination in a cell-based ubiquitination assay (**Fig. 1C**). We immunoprecipitated (IP) LMO7 to determine if it interacted with tau but found no evidence for tau-LMO7 interaction (**fig. S1, A)**. These results suggest that LMO7 regulates tau-ubiquitination indirectly. To identify potential candidates mediating this effect, we performed LMO7 IP followed by mass spectrometry (MS) analysis and identified 13 putative LMO7 interactors that are known to interact with tau (**Supplemental table 1**). Among these, the E3 ubiquitin-protein ligase CHIP emerged as a strong candidate mediator given its well-established role in tau ubiquitination and degradation^17–19^. We performed a Co-IP assay in cells that confirmed the LMO7-CHIP interaction (**Fig. 1D**).

Given the established role of CHIP in promoting clearance of abnormal tau and our observation that reducing LMO7 levels causes a decrease in tau, we hypothesized that LMO7 stabilizes tau by negatively regulating CHIP abundance or its E3-ligase activity. Interestingly, we found that LMO7 overexpression increased CHIP levels (**Fig. 1E-i**), while LMO7 knockdown decreased CHIP levels in cultured cells (**Fig. 1E-ii**). The observation that overexpression of LMO7 results in an increase of both CHIP and tau levels suggests that LMO7 inhibits CHIP ligase activity, which prevents its own autoubiquitination leading to an increase in CHIP levels^39,40^. Thus, LMO7 may inhibit CHIP’s enzymatic activity, rendering CHIP unable to ubiquitinate tau as well as itself.

To test whether LMO7 affects CHIP’s E3-ligase activity, we analyzed CHIP’s autoubiquitination in cells and found that LMO7 overexpression reduced CHIP polyubiquitination **(Fig. 1F),** confirming that LMO7 inhibits CHIP’s E3 ligase function.

Next, we examined whether LMO7 modulates CHIP-mediated tau-ubiquitination. Overexpressing LMO7 reduced tau-ubiquitination, whereas CHIP overexpression increased tau-ubiquitination. When LMO7 and CHIP were co-expressed, CHIP-induced tau-ubiquitination was completely abolished, resulting in ubiquitination comparable to when LMO7 is expressed alone (**Fig. 1G**). Together, these results demonstrate that LMO7 antagonizes CHIP by inhibiting its E3 ligase activity and thereby stabilizes both CHIP and tau proteins.

### LMO7 mediates interaction of CHIP with its inhibitor, BAG5

To understand the mechanism by which LMO7 inhibits the ligase activity of CHIP, we revisited our list of LMO7-interacting proteins. Among the candidates, we focused on the co-chaperone, BAG5, a known CHIP-binding partner that suppresses CHIP’s ubiquitin ligase activity^20,31,32^. Our co-immunoprecipitation (co-IP) data confirmed that BAG5 interacts with both LMO7 and CHIP (**Fig. 1H** and **1I**).

Given that LMO7 binds to both BAG5 and CHIP, we hypothesized that LMO7 might promote CHIP-BAG5 complex formation, thereby inhibiting CHIP activity. In support of this idea, co-IP followed by immunoblotting (IB) revealed that LMO7 knockdown decreased the CHIP-BAG5 interaction, as evidenced by the marked loss of CHIP co-IP in BAG5 IP compared with the control non-targeting (NT) shRNA (**Fig. 1J)**. Consistently, LMO7 overexpression enhanced the CHIP-BAG5 association (**Fig. 1K**). In contrast, reduction of BAG5 showed no effect on CHIP-LMO7 interaction (**fig. S1, B**). Similar to our cell-based findings, LMO7 knockdown also reduced the CHIP-BAG5 interaction in mouse brain tissue (**fig. S1, C**).

Together, these findings indicate that LMO7 stabilizes tau by promoting the CHIP–BAG5 interaction, which in turn suppresses CHIP’s E3 ligase activity and consequently limits tau ubiquitination and degradation.

### *Lmo7* knock-down rescues memory deficits and mitigates tau pathology in a tauopathy mouse model

To evaluate the therapeutic potential of targeting LMO7 in AD and other tauopathies, we examined whether partial reduction of Lmo7 could reverse pathological phenotypes in a tauopathy mouse model. We reduced *Lmo7* in adult P301S transgenic mice (PS19 line), using a doxycycline (Dox)-inducible shRNA expression system delivered via ICV injection at P0, as previously described^10^.

*Lmo7* knockdown was initiated at 2 months of age by switching from Dox-containing to regular food. After aging for 8 months, PS19 mice injected with NT-shRNA (NT-shRNA PS19) mice displayed reduced freezing behavior in the contextual fear conditioning task, compared to the age-matched NT-shRNA WT controls. In contrast, *Lmo7* knockdown rescued these learning and memory deficits: both WT and PS19 animals injected with *Lmo7* shRNA showed no difference in freezing compared to the NT-shRNA WT controls (**Fig. 2A**).

**Figure 2:**
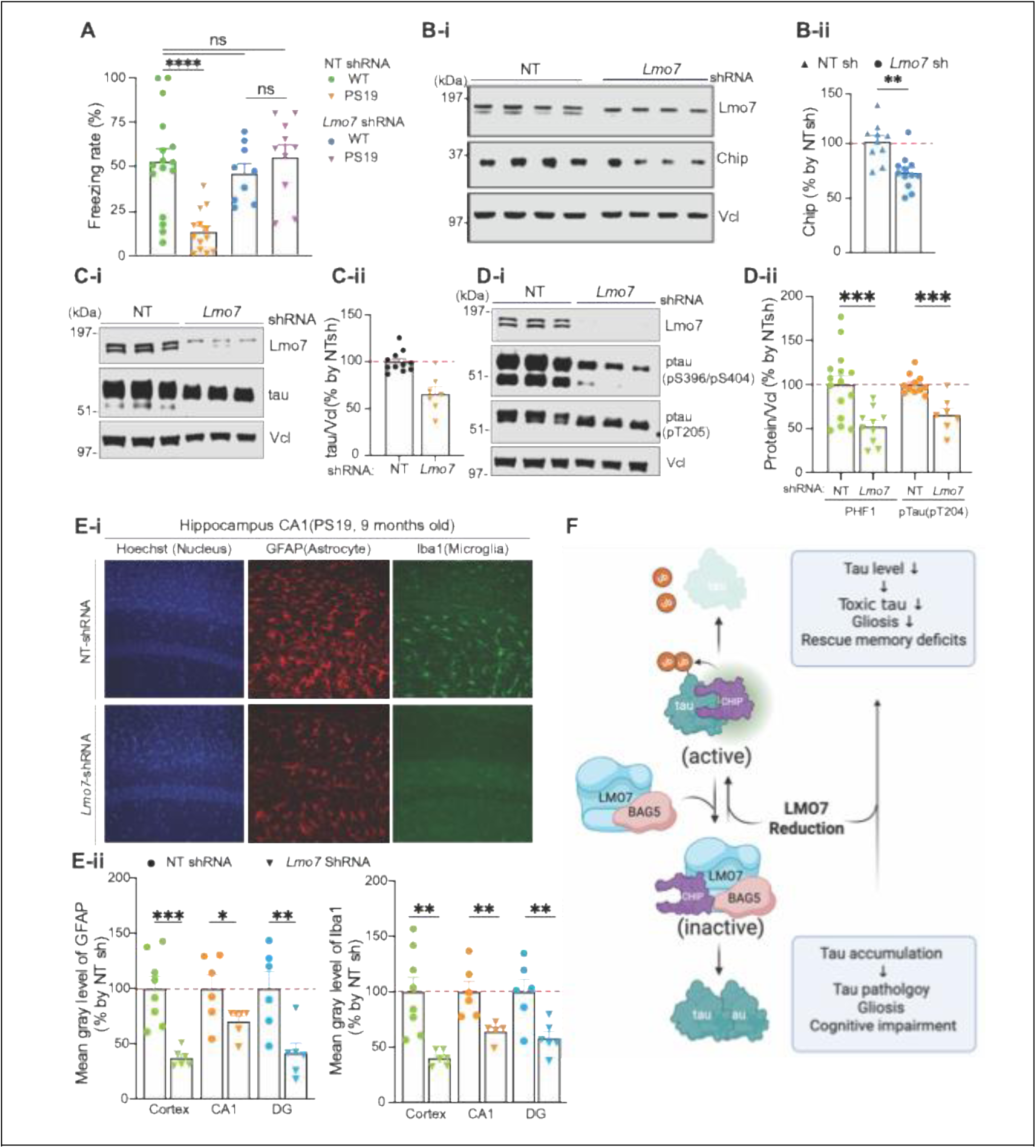
*Lmo7* knockdown rescues memory deficit and reduces pathogenic tau species and gliosis in the PS19 mouse brain. **(A)** Contextual fear-conditioning (CFC) task at 8 months in P301S tau transgenic mice (PS19) and non-transgenic littermate mice with or without knockdown of *Lmo7*. (**B-D**) IB images **(-i)** and quantification graph **(-ii)** of Chip **(B),** total tau **(C)**, and phospho-tau (**D;** pS396/S404 and pT205) in cortex and hippocampus from PS19 mice at 9 months of age, with or without *Lmo7* knockdown. **(E)** Neuroinflammation analysis in PS19 mice. **(E-i)** Representative images of astrocyte (GFAP) and microglia (Iba1) staining in the hippocampal CA1 region of 9-month-old PS19 mice expressing NT shRNA control or *Lmo7* shRNA. **(E-ii)** Quantification graphs of Iba1 (left) and GFAP (right) signal intensity in the cortex, CA1 region, and dentate gyrus (DG) in PS19 mice, with or without *Lmo7* knockdown. Each data point represents an individual animal, with an average of 2-3 sections analyzed per animal (n=5-8). **(F)** Schematic illustrating a potential mechanism by which LMO7 regulates tau levels. CHIP ubiquitinates tau and promotes its degradation. LMO7 antagonizes CHIP by promoting its interaction with inhibitory BAG5, leading to tau accumulation. LMO7 reduction enhances CHIP activity, reduces tau accumulation, and mitigates tau-associated toxicity. Data shown as mean ± SEM (*p≤0.05, **p≤0.01, ***p≤0.001, ****p≤0.0001).

Following behavioral testing, we evaluated the mouse brain tissues. First, we confirmed that *Lmo7* knockdown (KD) by shRNA reduced Lmo7 by 50-60 % (**Fig. 2B-i**). *Lmo7* KD reduced Chip protein levels by about 25 %, consistent with our findings in human cells and likely owing to its enhanced ubiquitination activity (**Fig. 2B**). Importantly, *Lmo7* KD markedly decreased total tau and pathological phospho-tau (pT205 and PHF [pS396/S404]) (**Fig. 2C** and **2D**). Lastly, we assessed gliosis in these PS19 mice. *Lmo7* KD reduced microgliosis, as evidenced by the decreased Iba1 immunoreactivity in the cortex, hippocampus CA1, and dentate gyrus (DG), compared to NT-shRNA PS19 mice. Similarly, abnormal astrocytic activation was attenuated, with reduced GFAP staining observed in the cortex, hippocampus and DG compared to control (**Fig. 2E**).

Collectively, our data demonstrate that LMO7 stabilizes tau by suppressing the E3 ubiquitin ligase activity of CHIP through enhancing its interaction with the inhibitory cochaperone BAG5. Partial reduction of Lmo7 enhances Chip E3 ligase activity. This in turn lowers tau levels, alleviates gliosis, and rescues cognitive deficits in tauopathy mice, highlighting LMO7 as a promising therapeutic target for tauopathies **(Fig. 2F)**.

### Dissociating the LMO7-CHIP interaction by CHIP-derived competing peptide activates CHIP and reduces tau accumulation

Together with the well-established neuroprotective role of CHIP, our findings suggest that disruption of the CHIP-LMO7-BAG5 complex represents a promising therapeutic approach to activate CHIP and prevent pathogenic accumulation of disease driving proteins such as tau. As a proof-of-concept, we examined whether interfering with the LMO7-CHIP interaction modulates CHIP activity and tau levels. Because no small molecule inhibitors of LMO7-CHIP binding are currently available, we designed a small CHIP-derived peptide corresponding to the LMO7-binding region to competitively disrupt their interaction. Using co-IP of LMO7 with a series of truncated CHIP constructs, we identified the TPR (tetratricopeptide repeat) and HH (helix-helix) domains of CHIP as essential for LMO7 binding (**Fig. 3A**). Integrating our experimental results with AlphaFold-based structural predictions of the CHIP–LMO7–BAG5 complex highlighted specific CHIP residues likely to mediate LMO7 binding (**Fig. 3B**). Based on these data, we selected residues 170–230 of CHIP as a candidate competing peptide (CHIP_170-230_).

**Figure 3:**
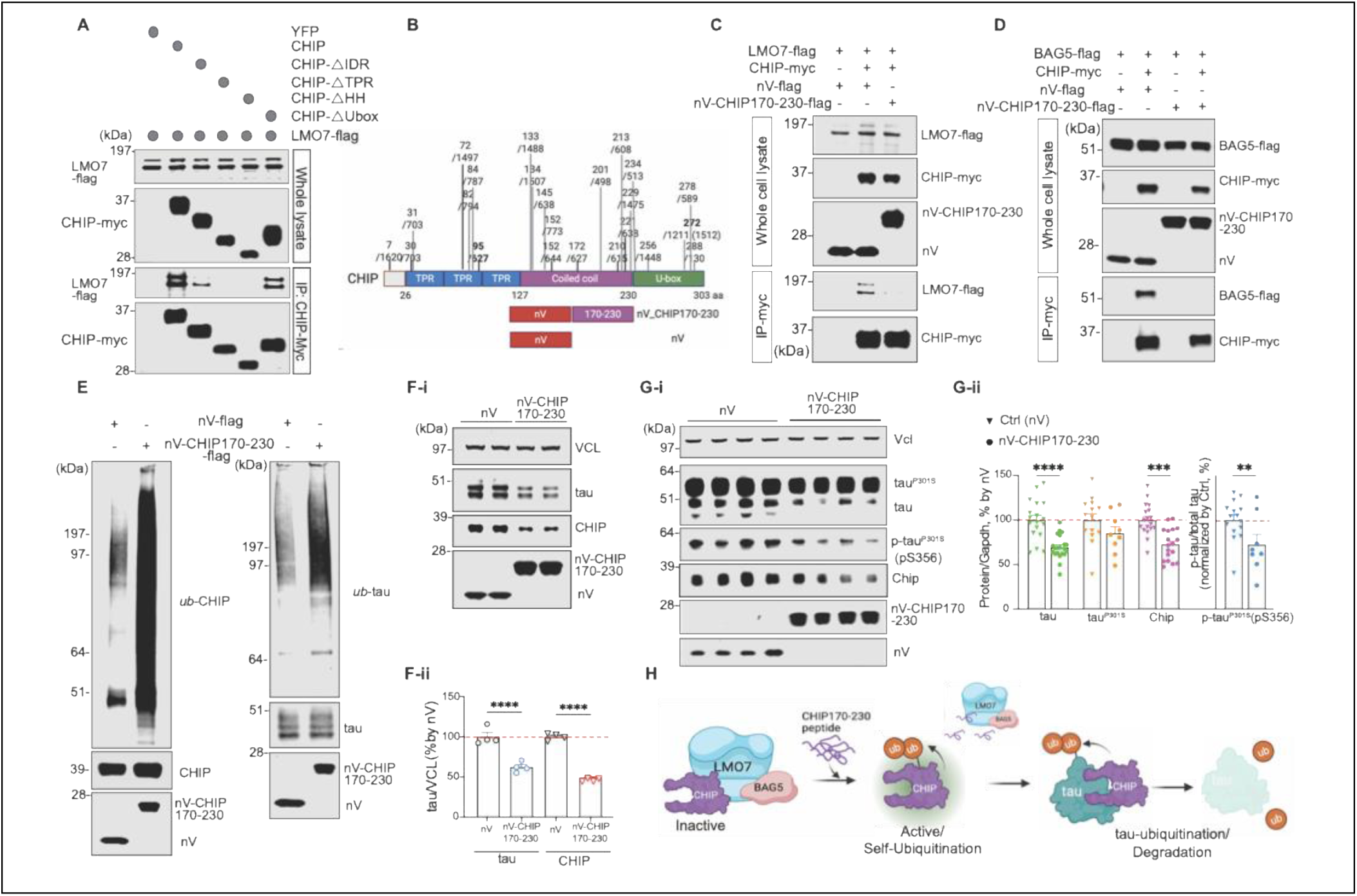
Disruption of LMO7-CHIP interaction by a CHIP-derived peptide reduces pathological tau species and demonstrates its therapeutic potential. **(A)** Co-IP of LMO7 by full-length or individual truncated CHIP constructs in HEK293T cells. **(B)** Location of the CHIP-derived peptide (CHIP_170-230 aa) that mediates interaction with LMO7, based on AlphaFold structural analysis of potential LMO7-interaction sites on CHIP. **(C)** IB image showing co-IP of LMO7 by CHIP IP in HEK293T cells overexpressing either control peptides or CHIP_170-230 peptide. **(D)** IB images showing Co-IP of BAG5 by CHIP IP in HEK293T cells overexpressing control peptide or CHIP_170-230 peptide. **(E)** Poly-ubiquitination of CHIP (left IB image) and tau (right IB image) in HEK293T cells overexpressing control peptide or CHIP_170-230 peptide. IB images **(F-i)** and quantification graph **(F-ii)** of decreased tau and CHIP protein in HEK293T cells by CHIP_170-230 aa peptide. **(G)** IB images **(G-i)** and quantitative graphs **(G-ii)** showing proteopathic mutant tau (tauP301S), endogenous wildtype mouse tau (tau), p-tau (pS356), and CHIP, in the tauopathy mouse model (PS19 line) overexpressing control peptide or CHIP_170-230 peptide. **(H)** Schematic diagram which show how inhibition of LMO7-CHIP interaction by CHIP_170-230 peptide restores CHIP’s activity and promotes tau’s clearance. Data shown as mean ± SEM (**p≤0.01, ***p≤0.001, ****p≤0.0001).

Overexpression of CHIP_170-230_ disrupted LMO7-CHIP interaction, as evidenced by the markedly reduced LMO7 co-IP in CHIP pull-down compared with the expression of a control peptide (**Fig. 3C**). Notably, disrupting LMO7-CHIP interaction also led to a loss of CHIP-BAG5 association (**Fig. 3D**). These data support our model that LMO7 mediates CHIP-BAG5 complex formation. Functionally, expression of CHIP_170-230_ enhanced CHIP’s E3 ligase activity, increasing polyubiquitination of both CHIP and tau (**Fig. 3E**), and consequently reduced CHIP and tau levels in cells (**Fig. 3F**).

Our previous studies, together with the data in this study demonstrate that even modest (10-20 %) reduction of proteopathic mutant tau is sufficient to ameliorate neurodegenerative phenotypes, including memory decline, in a tauopathy mouse model^7,8,10^. Building on this knowledge, we next evaluated whether inhibiting LMO7-CHIP interaction could reduce tau in a tauopathy mouse model. Viral overexpression of CHIP_170-230_ by AAV-ICV injection in the brain of PS19 tauopathy mice led to decreased levels of total tau and phosphorylated tau, as well as reduced CHIP abundance (>15 %) likely owing to the enhanced E3 ligase activity and auto-ubiquitination (**Fig. 3G**).

Collectively, these results demonstrate that disrupting the CHIP-LMO7-BAG5 complex, either with competitive peptides or potentially with small molecules, represents a viable strategy to activate CHIP and mitigate tauopathy-associated neurodegeneration (**Fig. 3H**).

### AI-driven identification of small-molecule inhibitors targeting LMO7-CHIP-BAG5 complex

Building on the therapeutic potential of disrupting the CHIP–LMO7–BAG5 complex, we employed an AI-driven drug repurposing platform (PharmacoNet)^38^ to identify FDA-approved compounds that might inhibit the formation of the CHIP-LMO7-BAG5 complex.

First, we defined the binding interface between LMO7 and CHIP (**Fig. 4A-i**) using AlphaFold-based structural predictions to identify sites for potential inhibition within the predicted complex. Each FDA-approved compound in the database was computationally evaluated for binding to this pocket (**Fig. 4A-ii**) through structure-based docking and affinity predictions. Compounds were then ranked according to their calculated binding energy (kcal/mol) and predicted binding affinity (µM) (**Supplementary Material 1**). From this screen, eight top-ranked candidates were experimentally tested. Six of these compounds significantly reduced both CHIP and tau protein levels in HEK293T cells (**Fig. 4B** and **4C**)

**Figure 4.**
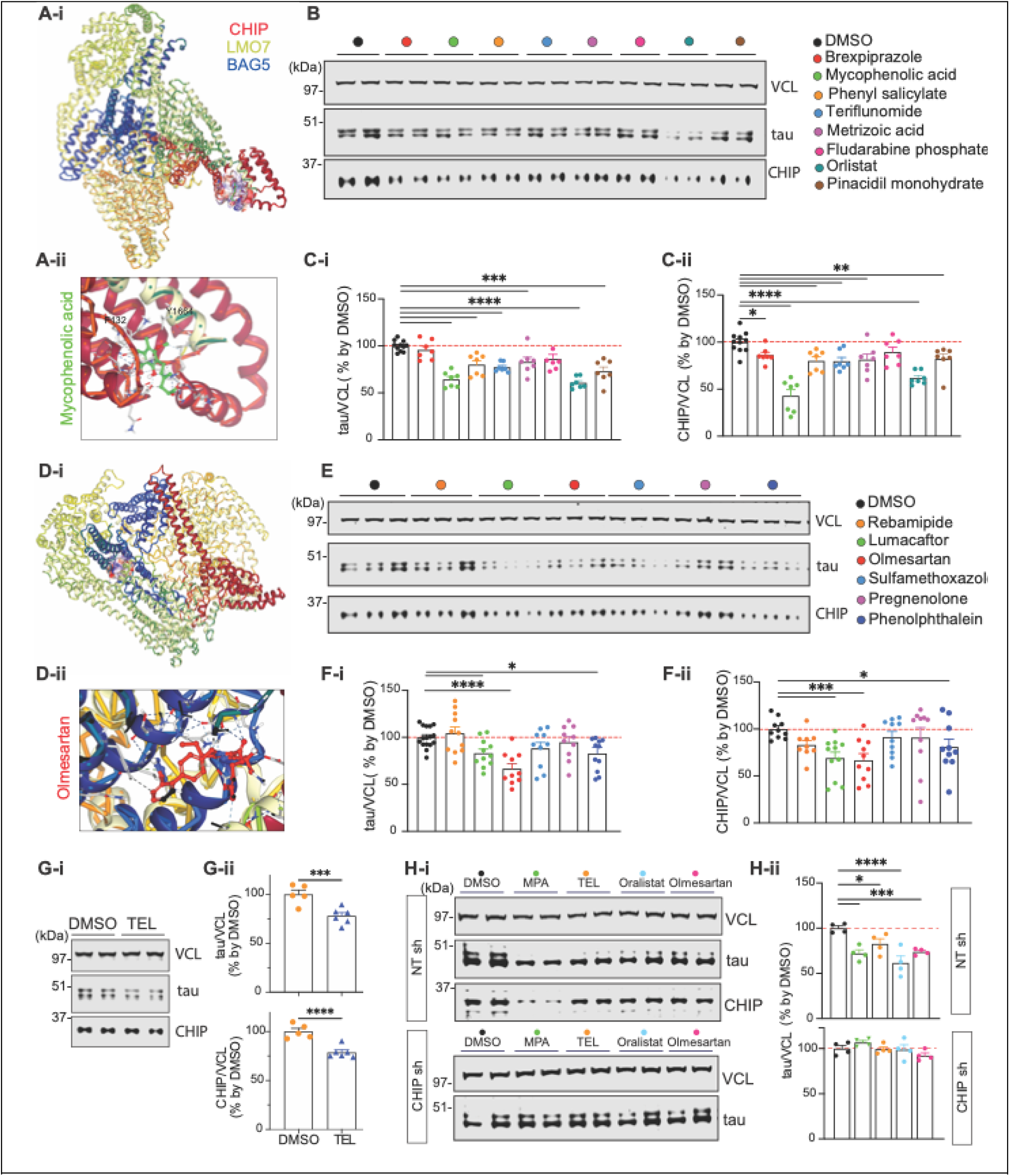
Identification of compounds that disrupt LMO7-CHIP or LMO7-BAG5 interaction using AI-based computational screening of FDA-approved medicines. (**A-i**) AlphaFold-predicted structure of CHIP-LMO7-BAG5 complex highlighting the potential interaction interface between CHIP and LMO7. **(A-ii)** A structural model showing Mycophenolic acid intercalating at the CHIP and LMO7 interaction interface. **(B)** Representative IB image of tau and CHIP in HEK293T cells treated with either vehicle control (DMSO) or top candidate compounds identified by screening targeting the LMO7-CHIP interaction. **(C)** Quantitative graphs of tau **(C-i)** and CHIP **(C-ii)** proteins in HEK293T cells treated with DMSO or the indicated compounds (1μM for 48 hours). **(D-i)** Structure prediction of CHIP-LMO7-BAG5 complex highlighting the potential interaction interface between LMO7 and BAG5. **(D-ii)** Structural model showing Olmesartan intercalating at the interface between LMO7 and BAG5. **(E)** Representative IB images of tau and CHIP in HEK293T cells treated with either DMSO or selected compounds identified from screening targeting LMO7-BAG5 interaction (1μM for 48 hours). **(F)** Quantitative graphs of tau **(F-i)** and CHIP **(F-ii)** protein levels in HEK293T cells treated with the indicated compounds. **(G)** IB image **(G-i)** and quantification graphs **(G-ii)** of tau and CHIP proteins levels in HEK293T cells treated with DMSO or Telmisartan (1μM for 48 hours). **(H)** IB image **(H-i)** and quantification graphs **(H-ii)** of tau protein levels in HEK293T cells treated with DMSO or the indicated compounds in the presence or absence of CHIP by NT shRNA or *CHIP* shRNA. Data shown as mean ± SEM (*p≤0.05, **p≤0.01, ***p≤0.001, ****p≤0.0001).

A similar computational pipeline was applied to identify inhibitors of the LMO7–BAG5 interaction (**Fig. 4D**). Following binding site definition and virtual screening, six top candidates were selected, of which three compounds significantly reduced CHIP and tau levels (**Fig. 4E** and **4F**).

Among the validated inhibitors, we selected Mycophenolic acid (MPA) and Olmesartan for further analysis. MPA is an immunosuppressive agent that exhibits partial blood-brain barrier (BBB) permeability. In contrast, Olmesartan is an angiotensin II receptor antagonist used to treat hypertension. Olmesartan lacks BBB penetration, making it less favorable for treating neurodegenerative diseases within the central nervous system.

Given that Olmesartan has several structural analogs that bind to the same target and share similar structural features, we evaluated Telmisartan (TEL) as an alternative inhibitor of the LMO7-BAG5 interaction, given its established BBB permeability. We demonstrated that TEL reduced CHIP and tau levels effectively in cultured cells (**Fig. 4G**). Both MPA and TEL displayed dosage-dependent effects on CHIP and tau levels, suggesting that their biological effect is specific and controllable rather than non-specific or cytotoxic (**fig S2, A** and **B**). We further confirmed that the identified compounds reduce tau in a CHIP-dependent manner, as their effect on tau levels was completely abolished in CHIP-deficient cells (**Fig. 4H**)

Mechanistically, MPA disrupted the CHIP-LMO7 interaction, resulting in loss of both LMO7 and BAG5 co-IP with CHIP. In contrast, TEL selectively disrupted the CHIP–BAG5 association while preserving CHIP–LMO7 binding, supporting our model and demonstrating target-specific inhibition by these compounds (**Fig. 5A**).

**Figure 5.**
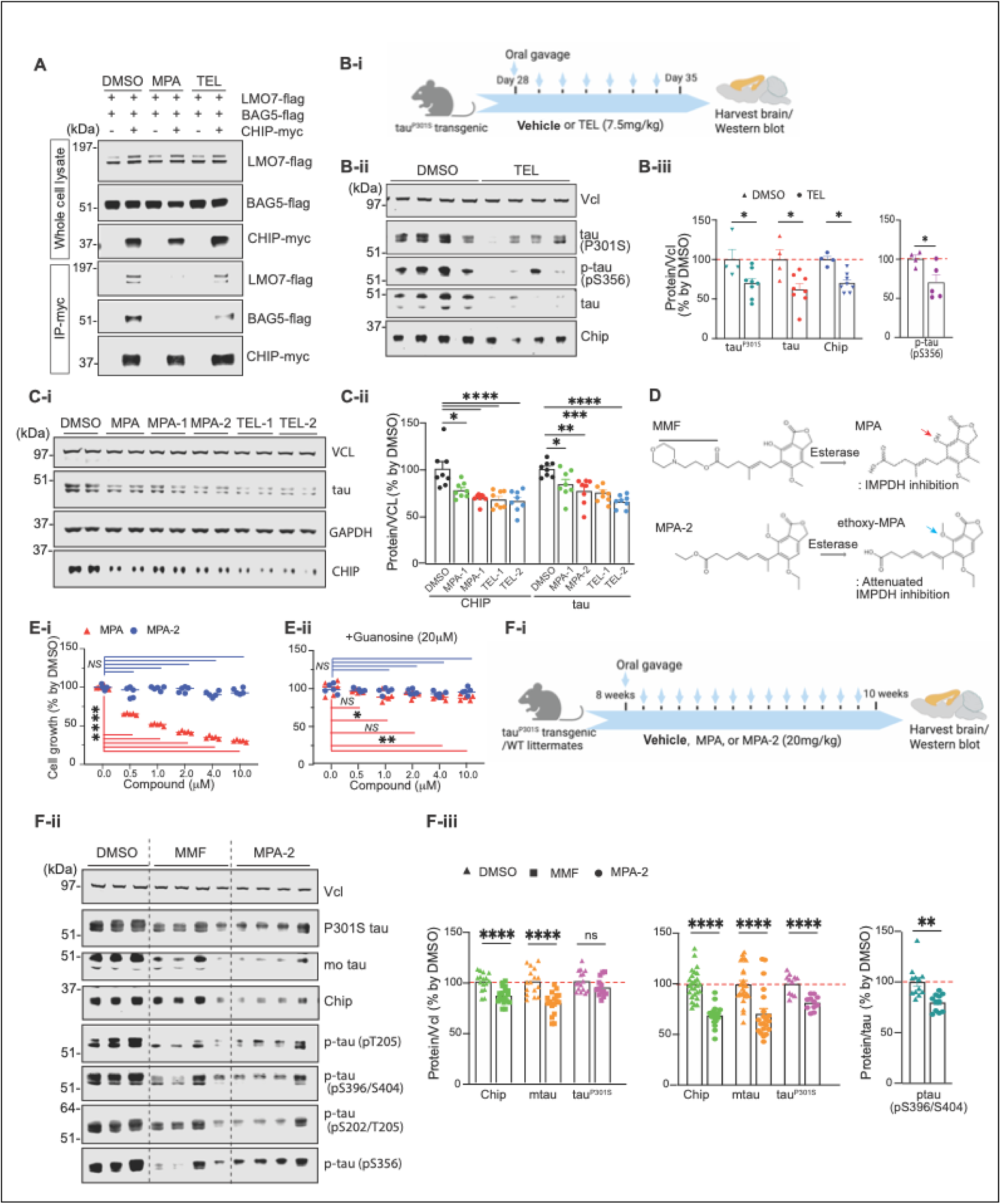
Telmisartan (TEL), Mycophenolic acid (MPA) and MPA analog (MPA-2) reduce pathological tau species in tauopathy mice brains. (**A**) IB image showing co-IP of LMO7 and BAG5 by CHIP-IP in HEK293T cells overexpressing LMO7-flag, BAG5-flag or /and CHIP-Myc. Cells were treated with DMSO, MPA (0.5μM), or TEL (1μM). **(B-i)** Schematic diagram of the experimental design testing TEL using oral gavage administration. Representative IB image **(B-ii)** and quantification graphs **(B-iii)** of tau, p-tau (pS356), and Chip in brain lysates from PS19 mice treated with either DMSO or TEL (7.5mg/kg, daily for 7 days). **(C)** Representative IB images (**C-i**) and quantification graph (**C-ii**) of tau and CHIP levels in HEK293T cells treated with MPA (0.25μM) or individual analog compounds (1μM). (**D**) Chemical structure of Mycophenolate mofetil (MMF, MPA prodrug), a modified MMF analog (MPA-2), and their esterase-cleaved products, MPA and ethoxy-MPA, highlighting different type of moiety at position 4 (arrow). (**E**) Growth curves of HEK293T cells treated with DMSO, the serial dilution MPA, or MPA-2, with or without restoration of intracellular GMP level by guanosine supplementation. **(F-i)** Schematic diagram of *in vivo* experimental design to test the efficacy of MMF (MPA prodrug) and MPA-2 using oral gavage administration. Representative IB image **(F-ii)** and quantification graphs **(F-iii)** of tau, p-tau, and Chip in brain lysate from tauopathy mice treated with DMSO, MMF or MPA-2 (20mg/kg, daily for 14 days). Data shown as mean ± SEM (*p≤0.05, **p≤0.01, ***p≤0.001, ****p≤0.0001).

### Oral administration of Telmisartan reduced pathologic tau species

Given that BBB penetration of TEL is better than that of MPA, we tested the *in vivo* efficacy of TEL using the PS19 tauopathy mouse model that we used to test the beneficial effect of LMO7 knockdown. Oral administration of TEL (**Fig. 5B-i**) significantly reduced both proteopathic tau and wild-type endogenous mouse tau as well as Chip in cortical tissue from PS19 mice, compared to vehicle treated mice. Moreover, brain tissue from TEL-treated mice also showed reduction in pathological phosphorylated tau (pS356) (**Fig. 5B-ii, -iii**). These data demonstrate that TEL provides a promising strategy to enhance CHIP activity and reduce tau protein prone to pathological aggregation, providing an opportunity to treat tauopathies.

### Structure-optimized MPA analogs activate CHIP and reduce tau, independent of inosine-5′-monophosphate dehydrogenase (IMPDH) inhibition

Next, we designed new molecules by modifying the original structure of MPA and TEL to improve blood-brain barrier penetration and enhance selectivity, biosafety, and bioavailability (**Supplementary Table 2**). MPA analogs (MPA-1 and MPA-2) and TEL analogs (TEL-1 and TEL-2) significantly reduced tau level in HEK293T cells (**Fig. 5C**). NanoBiT assay in live cells confirmed that the selected compounds inhibited CHIP-BAG5 interaction by disrupting LMO7-CHIP interaction (**fig. S2, C**). In addition, these compounds moderately but significantly enhanced tau-ubiquitination, as measured by NanoBiT-adapted mono-ubiquitination assay (**fig. S2, D**). Among them, MPA-2 and TEL-2 exhibited clear dosage-dependent effects on both CHIP and tau levels (**fig. S2, E** and **F**).

MPA is a well-known inhibitor of inosine-5′-monophosphate dehydrogenase (IMPDH), leading to guanine nucleotide depletion. To reduce IMPDH inhibition and avoid potential toxicity, MPA-2 was designed to diminish IMPDH binding. Derived from the MPA prodrug, Mycophenolate mofetil (MMF), MPA-2 is cleaved by esterase to release the ethoxy-modified MPA in which C4 hydroxyl group critical for IMPDH interaction is altered, resulting in markedly reduced intrinsic affinity for IMPDH while preserving its tau-lowering activity (**Fig. 5D**).

At higher concentrations (∼1 μM), the original compounds MMF or MPA further enhanced tau reduction without affecting pS6, pAMPK, or p53 levels (**fig. S3, A**). However, they decreased protein abundance (**fig. S3, B**), and cell proliferation (**Fig. 5E**), indicating guanine nucleotide–limited cytostasis independent of canonical stress or energy pathways. In contrast, MPA-2 lowered tau without altering protein synthesis, cell proliferation, or stress markers, indicating lack of IMPDH inhibition and a safer mode of CHIP activation. Guanosine supplementation restored MPA-induced protein-synthesis and growth suppression via the salvage pathway but had no effect on tau reduction by MPA or MPA-2, producing only partial attenuation for MPA (**fig. S3, C**). Collectively, these findings demonstrate that our newly designed molecules lowered tau independent of nucleotide depletion or global translational inhibition. Notably, our data show that the newly designed MPA-2 disrupts CHIP-LMO7 interaction and reduces tau levels without IMPDH inhibition and accordingly its associated toxicity.

### MPA and MPA analog reduced pathologic tau species

Oral administration of MPA for 14 days (**Fig. 5F-i**) significantly reduced Chip and endogenous wild-type mouse tau in cortical tissue from PS19 mice, but it did not appreciably lower exogenous human pathogenic tau, or its disease-associated phosphorylated species when compared with vehicle-treated controls (**Fig. 5F-ii, -iii upper graph**). In contrast, the same regimen of MPA-2 robustly decreased both endogenous and proteopathic tau species as well as Chip, demonstrating superior *in vivo* efficacy (**Fig. 5F-ii, -iii**).

Together, these findings demonstrate that pharmacological activation of CHIP via MPA and our newly designed derivative compound represents a promising and well-tolerated new strategy to treat Alzheimer’s disease and other tauopathies *in vivo*.

## Discussion

Here we identify the CHIP–LMO7–BAG5 complex as a previously unrecognized control point for tau homeostasis and a tractable therapeutic target for tauopathies. Three findings make this work distinctive. First, we resolve a long-standing challenge in the field by defining a mechanism to indirectly activate CHIP, a broadly protective E3 ubiquitin ligase implicated in multiple neurodegenerative diseases, through disruption of its endogenous inhibitory complex. Second, we show that this complex can be disabled *in vivo* — genetically, with a competing peptide, or with orally available small molecules — to reactivate endogenous CHIP, lower tau, attenuate gliosis, and rescue cognition. Third, by integrating cross-species genetics, AlphaFold-guided interface design, and AI-based virtual screening of FDA-approved drugs, we converted a screen hit into *in vivo*-active leads (Telmisartan, MMF/MPA, MPA-2). To our knowledge, this is the first demonstration that an FDA-approved drug repositioned through AI-based docking lowers tau in a mammalian tauopathy model.

Mechanistically, LMO7 acts as a scaffold that bridges CHIP and BAG5, suppressing CHIP’s E3 ligase activity and limiting the degradation of tau. We also identified a small peptide and small-molecule inhibitors that enhance CHIP activity by disrupting the CHIP-LMO7-BAG5 complex. Disrupting this complex lowers the level of total tau and pathogenic hyperphosphorylated tau species in a tauopathy mouse model. Importantly, optimization of the lead compound MPA yielded MPA-2 an analog that selectively enhances CHIP activity without IMPDH inhibition or cytotoxicity, demonstrating the feasibility of a safe and targeted therapeutic strategy.

Although BAG5 has been reported to inhibit CHIP^20,31,32^, how BAG5 engages CHIP has remained unresolved for nearly two decades. We show that CHIP does not bind to BAG5 directly; instead, LMO7 bridges this interaction, as disrupting CHIP-LMO7 binding with CHIP_170-230_ or small molecules abolishes CHIP-BAG5 binding. LMO7 thus functions as a previously unrecognized scaffold that permits BAG5 to act on CHIP, transforming a binary cofactor relationship into a tractable trimeric complex. Future *in vitro* binding assays and high-resolution structural studies of the CHIP–LMO7–BAG5 complex by cryo-EM will further define the molecular architecture of this regulatory complex and enable structure-guided drug design.

Conceptually, scaffold-mediated E3-ligase reactivation is the inverse of targeted-degrader strategies (PROTACs, molecular glues), which couple a substrate to an active ligase^29^. Direct pharmacological activation of E3 ligases has been considered intractable because activation requires altering, rather than blocking, an enzymatic surface. Our results suggest a generalizable solution: where an inhibitory scaffold or cofactor exists, blocking the scaffold restores ligase activity using the cell’s own machinery. This logic should extend to other E3 ligases held in check by inhibitory cochaperones (e.g., Parkin–BAG5^41^, MDM2^32^ and partners in cancer biology) and identifies a broader class of druggable nodes in protein quality control.

Our group has shown that a modest reduction in tau – regardless of the upstream regulator, including kinases (Nuak1^8^ and TYK2^7^), E3-ligases (RNF130 and RNF149^10^), and ubiquitin hydrolases USP7^10^—produce *in vivo* benefits. Together with our previous findings, these data highlight the benefit of moderately lowering tau as a viable therapeutic strategy for AD and other tauopathies.

Adult brain-specific *Lmo7* knockdown in mice was well-tolerated and naturally occurring loss-of-function LMO7 variants are present in healthy individuals^42^, supporting the notion that partial inhibition of LMO7 is both safe and druggable. Importantly, our small-molecule strategy does not eliminate LMO7; it disrupts a single interaction interface, preserving LMO7’s other physiological functions while efficiently enhancing CHIP activity, and minimizing the unintended consequences of global LMO7 inhibition. The MPA-2 analog further decouples tau-lowering activity from IMPDH inhibition, suggesting a wide therapeutic index.

Previous studies have also shown that CHIP overexpression or pharmacologic activation can ameliorate diverse disease phenotypes in mice without overt toxicity^43^, further reinforcing the translational potential of targeting this pathway. Nevertheless, comprehensive physiological, behavioral, and toxicological studies will be required to fully evaluate long-term safety, pharmacokinetics, and efficacy of the molecules identified in this study prior to clinical application. That the chemical starting points are FDA-approved drugs with decades of clinical use, will de-risk the path forward.

Our findings are particularly compelling because, beyond tau-driven diseases, CHIP plays a protective role in multiple neurodegenerative diseases by preventing the accumulation and aggregation of pathogenic proteins such as ATXN1 in SCA1^21^ and α-synuclein in PD^20^. Moreover, loss of function variants in *STUB1* cause hereditary ataxias, including the recessive spinocerebellar ataxia 16 (SCAR16)^22,26,27^ and the *STUB1*-haploinsufficient SCA48^28^. Therefore, strategies that enhance CHIP activity hold substantial translational promise across both gain-of-function proteinopathies and loss-of-function CHIP disorders. Because direct pharmacological activation of E3 ligases remains challenging, our findings suggest that disrupting the CHIP-LMO7-BAG5 complex may provide a viable treatment approach for a range of neurodegenerative diseases.

In sum, we revealed a previously unrecognized role of LMO7 in antagonizing CHIP E3 ligase activity by promoting CHIP–BAG5 complex formation, limiting CHIP activity and tau degradation. Genetic or pharmacological disruption of this complex using competitive peptides, or MPA/TEL analogs activates CHIP, enhances tau clearance, and reduces pathogenic tau *in vivo*, with MPA-2 producing potent effects without IMPDH inhibition or cytotoxicity. These novel pharmacological modulators pave a path for the treatment of Alzheimer’s disease, other tauopathies, the *STUB1* ataxias, and other CHIP-associated neurodegenerative disorders. By integrating mechanistic genetics, AI-guided structural modeling, and drug repositioning, we provide a generalizable blueprint for reactivating endogenous E3 ligases held in check by inhibitory scaffolds.

## Materials and Methods

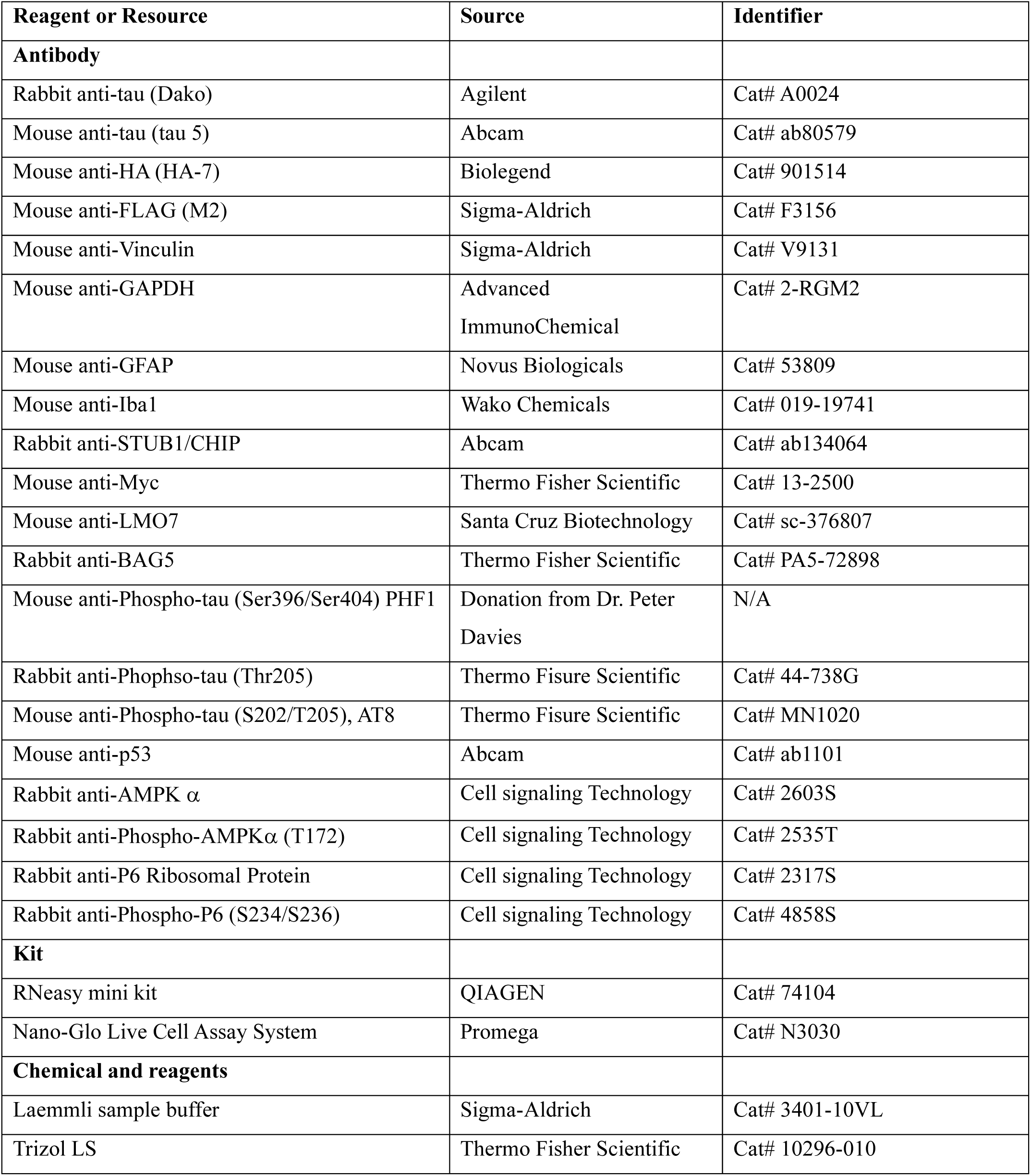
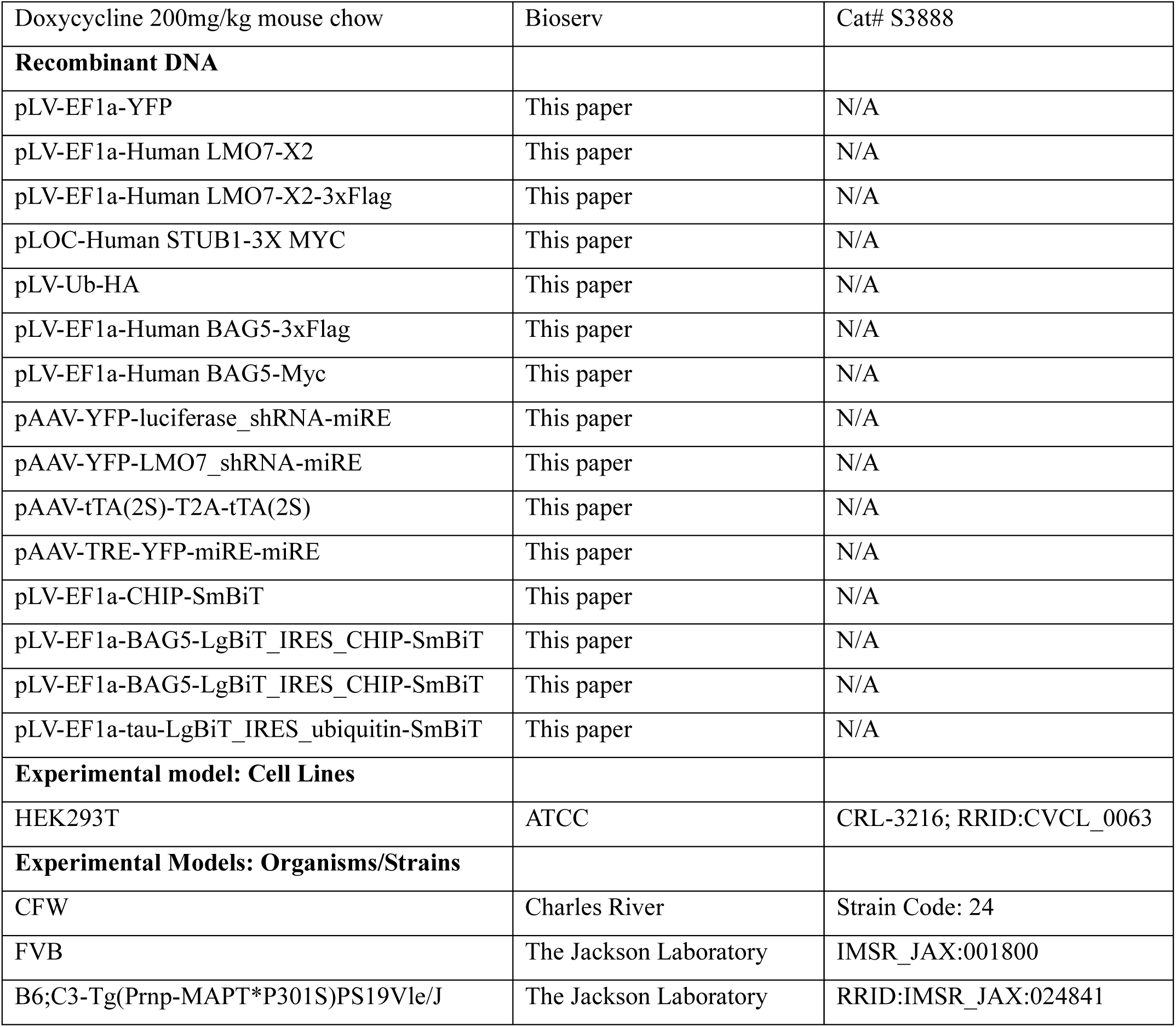

### Cloning

We designed 4-5 individual shRNA sequences for *Lmo7*, using the SplashRNA algorithm (REF70) and synthetized by IDT (https://www.idtdna.com/). shRNA sequence was amplified by PCR, using one pair of oligomer set (miRE-Gib-forward; 5’-ttcttaacccaacagaaggctcgagaaggtatattgctgttgacagtgagcg, miRE-Gib-reverse: 5’-gtaaacaagataattgctcgaattctagccccttgaagtccgaggcagtaggca), followed by integration into pAAV-YFP-miRE vector linearized by Xho1/EcoR1, through Gibson cloning (New England Biolab, E2621) as previously described^44^. The two independent shRNA that target *Lmo7* or Luciferase gene (non-targeting, NT) were integrated into the double miRE cassette containing tetracycline responsive element (TRE) one by one using same procedure described above for inducible expression.

Human *LMO7* splice variant 5 was purchased from DNASU (HsCD00871834), was amplified with 3X Flag tag at the C-terminal by PCR. cDNA encoding *LMO7* the first 285 N-terminal amino acids was synthesized by IDT and then cloned with *LMO7* spliced variant 5 into the linearized pLenti-EF1a (Addgene, #85132) cut by BamHI/EcoRI using a Gibson Assembly reaction. The resulting cDNA encodes LMO7 splice variant 2 which the neural tissue expressed isoform^33^.

Human full length *CHIP* plasmid was previously described (REF9). Four truncated CHIP (ΔHH, ΔIDR, ΔTPR and ΔUbox domains) were amplified from Human full length *CHIP* plasmid. PCR products were inserted into the linearized pLoc-DsRed plasmid cut by BamHI and NheI using Gibson assembly reaction.

### Viral production

Low-passage 293T cells at 80-90 % confluency was transfected with a 4:3:1 ratio of viral shuttle vector (pGIPz, pLenti, pLOC), psPAX2, and pMD2.G using Lipofectamine 2000 (Invitrogen, 11668019) or TransIT®-293 (Mirus, MIR2704) transfection reagent for lentiviral generation. We performed transfection according to manufacturer’s instructions. At 6-12 hours post-transfection, medium was replaced and then collected at 48- and 72-hours post-transfection. The viral suspension was then frozen and kept at −80 C ° until needed or concentrated 50-fold using Lenti-X concentrator (Clontech, 631231) when higher titer and more purified virus was required. Recombinant AAV8 was produced using a triple transfection with pAAV-, capsid-, and helper plasmid and the resultant virus was purified and concentrated on an iodixanol step gradient, as previously described^45,46^. To increase the yield of virus particles, cell-associated and medium-containing secreted AAVs were collected separately at 72 hours post-transfection and combined before purification for P0-ICV injection. For *in vitro* applications, the collected medium was concentrated 50-fold using AAVanced™ Concentration Reagent (SBI, AAV100A-1).

### *In vivo* gene delivery

The knockdown efficiency of *Lmo7* shRNA in pAAV was tested in Neuro2A cell as previously described^47^. The two most potent shRNAs targeting each gene were packed into AAV serotype 8 (AAV8). AAV8 harboring a shRNA expression cassette was bilaterally infused into the mouse brain at 4-10 × 10^10^ viral genomes per hemisphere using intracerebroventricular (ICV) injection targeting lateral ventricle at post-natal day 0 (P0) as previously described^48,49^. Transgenic offspring were generated by mating PS19 males with FVB females; pups were injected with the mixture of AAV8 containing either TRE-YFP- *Lmo7* shRNA - *Lmo7* shRNA or TRE-YFP- Luci shRNA-Luci shRNA in miRE backbone (8 × 10^10^ viral particles/hemisphere) and the TA2S-T2A-tTA2S expression cassette (2.0 × 10^10^ viral particles/hemisphere) using ICV injection at P0. Injected PS19 mice and WT littermates were fed with a doxycycline (DOX)-containing diet until 2 months post-injection. At 2 months, mice were switched to a DOX-free diet to allow for adult expression of the respective shRNA until 9 months of age for behavioral and pathology assessments.

### Contextual fear conditioning

The contextual fear conditioning test was performed as described in our previous study^10^. On training day, mice were habituated in home cages placed in behavioral room for 30 min. The individual mice were then placed in a soundproof operation box (20 cm length x 20 cm width x 35 cm height) and explored freely for 2 minutes. Three consecutive electrical foot-shocks were delivered to the mice at 1.5 mA for 2 seconds with 1-minute intervals. Mice were then returned to their home cage. The next day, we placed the individual trained mice and measured their freezing response in the same operation box for 5 minutes. The freezing response was scored as the total time mice spent ‘frozen’ during the test period. Data acquisition, control of stimuli, and data analysis were performed automatically using the Coulbourn/Actimetrics FreezeFrame3 System.

### Mouse brain sample preparation

For acute brain harvest from 3-week-old mice, mice were sacrificed by isoflurane inhalation. The hippocampus and caudal subregion of cortex above the hippocampus was dissected out and frozen on dry ice immediately. 9-month-old PS19 mice and age-matched WT littermate mice were sacrificed by sodium pentobarbital overdose and transcardially perfused with ice-cold PBS (20 ml per 25 g mouse) by gravity at 2.5 ml/min speed. Brains were isolated and split into two hemispheres. Cortex and hippocampus from the hemisphere were dissected out and frozen together. The remaining hemisphere was fixed by 4% paraformaldehyde (PFA) for 48 hours and cryopreserved in 1X PBS containing 30% sucrose at 4 °C until the samples sunk. Frozen samples were homogenized using an electric pestle (handheld polytron, WPR, 47747-370) in 10x volumes/weight of cold PEPI buffer (1x PBS containing 5 mM EDTA, protease inhibitor cocktail and phosphatase inhibitor cocktail). Part of the homogenate was then diluted to 1:1 in RIPA buffer (1x PPS containing 5 mM EDTA, protease inhibitor cocktail, phosphatase inhibitor cocktail, 1% deoxycholate, 1% Triton X-100, and 1% SDS) and gently mixed. After centrifugation at 15,000 RPM for 20 min, the supernatant was collected.

### Immunofluorescence and microscopy assay

Fixed brains were frozen on dry ice and sagittally sectioned (40 µm thickness) using a sliding microtome (Leica, SM2010 R). 3-5 sagittal sections per individual animal, containing hippocampus, lateral ventricle, and striatum (located 1-1.5 mm form the midline) were selected for staining. Brain sections were immunostained with GFAP (1:2,000; Novus Biologicals, 53809), and Iba1(1:2,000; Wako Chemicals, 019-19741) antibodies diluted in 1X PBS containing 0.25 % Triton X-100 and 7 % donkey serum at 4 °C overnight. Sections were then incubated in Alexa Fluor® 555 conjugated goat anti-rabbit IgG secondary antibodies (Invitrogen, A32732), Alexa Fluor® 647 conjugated donkey anti-mouse IgG secondary antibodies (Jackson ImmunoResearch, 715-605-150), and Hoechst (1:2,000, Abcam, ab228551) diluted in same buffer for 2 hours at RT. Fluorescent images were collected from nonoverlapping fields within the somatosensory cortex (above CA1), hippocampus CA1, and the dentate gyrus. A single optical plane of 0.977 µm in depth was collected at 405nm (Hoechst), 555nm (GFAP), and 647nm (Iba1) channels using fluorescent microscopy (Carl Zeiss) at 100 X magnification (895.26 µm x 670.8 µm per field). For representative images, Z-stacked fluorescent images (14 optical images) were collected from the same field of the hippocampus CA1 region using confocal microscopy (Carl Zeiss) at 200 X magnification (416.80 µm x 416.80 µm).

### Cell culture, transduction, infection, and drug treatment

Cells were transfected with plasmids encoding cDNA or short hairpin RNA using Lipofectamine 2000 (Invitrogen,11668019) or TransIT®-293 (Mirus, MIR2704) transfection reagent, according to the manufacturer’s protocol, and incubated for 48-72 hours followed by cell lysate preparation. Cells were transduced with lentivirus expressing ORF at 5-10 MOI in the various cell culture size formats. After 24 hours of transduction, cells were incubated with the appropriate antibiotics at least for 3 days. Cells were further maintained in antibiotics-diluted medium until additional treatment or harvest. At the end of the experiment, cells were washed with 1× PBS and frozen at −80 C° until further use.

### Cell lysate preparation

The cells in 24 well plate or 6 well plate-format were lysed by gentle mixing with ice-cold lysis buffer (50 mM Tris,150 mM NaCl, 2 mM CaCl2, 1 % Triton X-100, 5 mM EDTA, 5 % glycerol, pH7.5) containing 1x protease and phosphatase inhibitor cocktails (Genedepot, P3100-100, P3200-020) on shaker for 30-60 min at 4 °C. The whole cell lysate was ruptured by pipetting and then transferred to microfuge tubes. The supernatant was collected by centrifugation at 15,000 rpm for 30 min at 4 °C and then subjected to immunoprecipitation experiment or sample preparation by adding Laemmli sample buffer and denaturing (75 °C for 10 min) for immunoblotting experiments. Total protein concentration of lysate was determined by Bicinchoninic acid (BCA) assay using a BCA Protein Assay kit (10-009-D).

### Immunoprecipitation

Supernatant from one well of a 6-well plate was incubated with 0.75µg of tau antibody (Abcam, ab80579), 1 µg of Myc antibody (Thermo Fisher Scientific, 12-2500), or 1 µg of Flag antibody (Sigma, F3156) conjugated with protein G conjugated Dynabead (20 µl slurry per 1 µg antibody, Fisher Scientific, 10-009-D) for 2-4 hours at 4 °C. For ubiquitination assay supernatant was incubated with antibody for overnight. After washing with lysis buffer, 4-6 times, immunoprecipitated proteins were eluted in 35 µl of 1.5 x Laemmli sample buffer (Sigma-Aldrich, S3401-10VL) by boiling 75°C for 10 min. Brain lysates were incubated with 2ug of CHIP antibody (Abcam, ab134064) conjugated with protein A conjugated Dynabead (20 µl slurry per 1µg antibody, Fisher Scientific, 10002D) for 2-4 hours at 4°C.

### Western blot analysis

Cell or brain tissue lysate (1-5 μg/μl) in 1x sample buffer was denatured at 75° C for 10 min. To detect total tau from the cultured cells, total 20 μg of sample was incubated with α-tau antibody (Dako, 1:8000) whereas 5-10 μg (WT mouse) or 2-5 μg of sample (PS19 tau transgenic mouse) from mouse brain tissue was detected by α-tau antibody (Abcam, ab80579, 1:4000). Protein samples were resolved on a precast 4–12% Bis–Tris gels (MOPS running; Boston BioProducts, BP-178) and transferred onto nitrocellulose membranes (Biorad, 1620145) in Tris-glycine buffer supplemented with 10% methanol at 120 mV for 100 min at 4°C. After blocking 20 min with 5% bovine serum albumin (BSA), the membrane was incubated with the indicated primary antibodies overnight at 4° C. The protein was then visualized using corresponding fluorescent secondary antibodies (Odyssey, 1:10,000-20,000). Infrared fluorescence was measured with the Odyssey CLx imager (LI-COR) and quantified using Image Studio software (Li-COR Biosciences).

### IP and mass spectrometry

We overexpressed YFP (transient transfection control) and LMO7-3X Flag in HEK293T cells. After 48 hours, our transfection samples (∼2 × 10^8^ cells per sample) were collected and lysed as described above (“Cell lysate preparation”). Cell supernatant then incubated with Flag antibody conjugated agarose beads (Sigma-Aldrich, A2220) at 1μl bead for 2 million cells for 2 hours at 4 °C. After stringent washing with the same lysis buffer, IP samples were eluted by adding Laemmli sample buffer and denaturing (95 °C for 5 min). IP samples were digested with a gel after separation by SDS-PAGE to break down into smaller peptides for analysis by LC-MS/MS, using an AB SCIEX TripleTOF 5600 mass spectrometer. The data will be searched against the NCBI RefSeq database using Mascot, MS1-level analysis in Proteome Discoverer, and gene-centric identification and quantification with gpGrouper (REF PMID: 30093420). Output will include peptide MS1 AUC values and iBAQ-based protein quantification. For statistical reliability, we studied three replicates for each group by independent IP/MS experiments.

### In silico screening

The in silico screening workflow was performed using the Pharmaco-Net AI-driven drug discovery platform (https://pharmaco-net.org). Three-dimensional structure of human CHIP-LMO7 (isoform 2)-BAG5 complex was predicted using AlphaFold. Using PocketFinder module, we automatically identified a potential ligand binding site at either LMO7-CHIP or LMO7-BAG5 interface. This module applies an internal geometry- and energy-based algorithm to detect binding pockets while correcting common structural issues such as missing residues or alternative conformations. Following pocket identification, we selected a virtual screening library from the built-in FDA-approved compound collection (850 molecules, Enamine). For each compound, structure-based docking and binding affinity prediction were conducted using AI-AI-Dock and DeepCalici-plus modules by calculating a binding energy (kcal/mol) and dissociation constant (K_d_, µM). Candidate compounds were then ranked according to their predicted binding affinity and binding energy.

### Design and synthesis of analogs

New chemical entities (NCEs) were rationally designed using Mycophenolic acid (MPA), Telmisartan (TEL), and Olmesartan as backbone scaffolds identified from AI-based virtual screening of an FDA-approved drug library targeting the LMO7–CHIP–BAG5 complex. Structure–activity relationship analysis and medicinal chemistry optimization, supported by multiple AI models (Gemini, Grok, and ChatGPT), guided the design of analogs. Physicochemical properties were evaluated using RDKit and filtered by Lipinski and Veber rules, followed by virtual ADME and toxicity screening to prioritize CNS-suitable candidates. Selected compounds were synthesized by WuXi AppTech (Hong Kong).

### Oral gavage of small molecules

Each small molecules were weighed and dissolved in dimethyl sulfoxide to generate a stock solution. The stocks were freshly diluted in corn oil at a ratio of 1:17 (v/v) to yield the dosing formulation. 2-month-old tau P301S transgenic mice (PS19 line, [C57BL/6 x C3H] F1) received Telmisartan (7.5 mg/kg), MPA (20 mg/kg), and MPA-2 (20 mg/kg) or an equivalent volume of vehicle by oral gavage daily for one week for Telmisartan or two consecutive weeks for MPA and MPA-2. Dosing volumes were adjusted on each day according to body weights to ensure accurate drug delivery.

### Statistical Analyses

All statistical analyses were done using GraphPad Prism 9.0. Comparisons between two groups were analyzed using Student’s t test. Comparisons involving more than two groups were analyzed using one-way or two-way ANOVA followed by Bonferroni post hoc tests. All reported p-values are for post hoc comparisons. All graphs display group mean ± SEM (∗p ≤ 0.05, ∗∗p ≤ 0.01, ∗∗∗p ≤ 0.001, and ∗∗∗∗p ≤ 0.0001).

## AUTHOR CONTRIBUTIONS

J.K, B.T and H.Y.Z conceived the study, designed experiment, analyzed, and interpreted the data, and wrote manuscript. B.V, X.D, and Y.W performed cellular, molecular, and biochemical experiments. S.Y.J performed mass spectrometry and analyzed the resulting data. J.X designed the structurally modified analogs from the selected compounds.

## ACKNOWLEDGMENTS

We are thankful to the members of H.Y.Z laboratory for all the support, critical discussions and feedback that refined this manuscript. We thank Dr. Virginia Lee, University of Pennsylvania to generously share the PS19 mice. This research was supported by the Freedom Together Foundation, the Ting Tsung and Wei Fong Chao Foundation, the Hamill foundation, 5R21AG088501-02, and the Howard Hughes Medical Institute. We also would like to thank the Gene Vector Core of Baylor College of Medicine and the Neurobehavioral Cores at the Jan and Dan Duncan Neurological Research Institute (NRI) at Texas Children’s Hospital and the BCM-IDDRC (P50HD103555).

## Competing interests

H.Y.Z cofounded Cajal Neuroscience, is a Director of Regeneron Pharmceuticals board and is on the scientific advisory board of Cajal Neuroscience and the Column Group.

---

**Supplementary Table 1.**
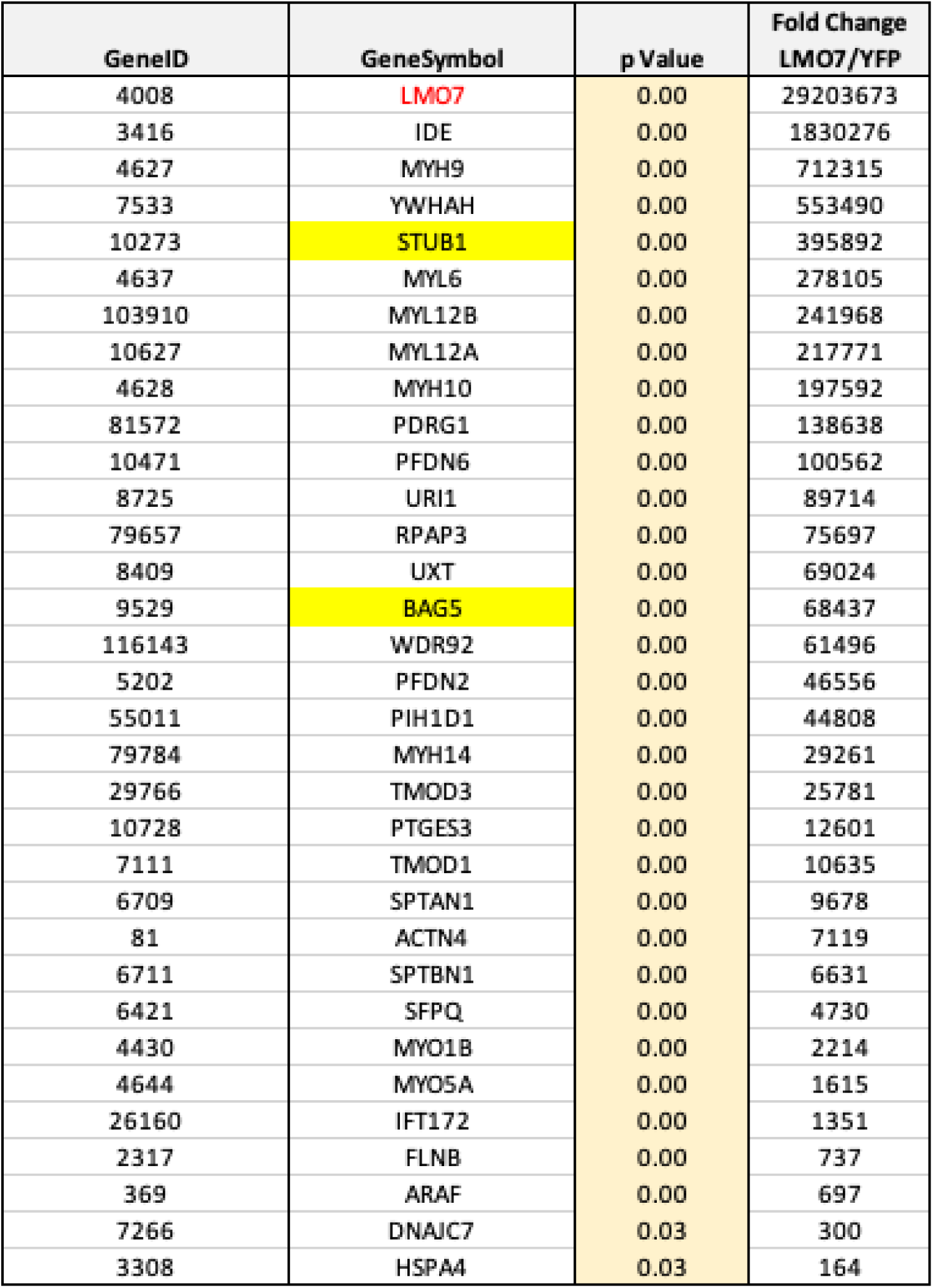
Results of IP-MS showing the proteins that co-precipitated with LMO7. Fold change indicates the abundance of a specific protein in the LMO7-IP sample compared to YFP control samples. Possible interactors are selected using the cut-off of Fold change >100 and significance p-value<0.05. LMO7 (in red) was efficiently pulled down via our IP and identified in our experimental samples but not in our control YFP. Both STUB1 (CHIP) and BAG5, highlighted in yellow under the Gene Symbol column, were identified as possible interactors of LMO7.

**Supplementary Table 2.**
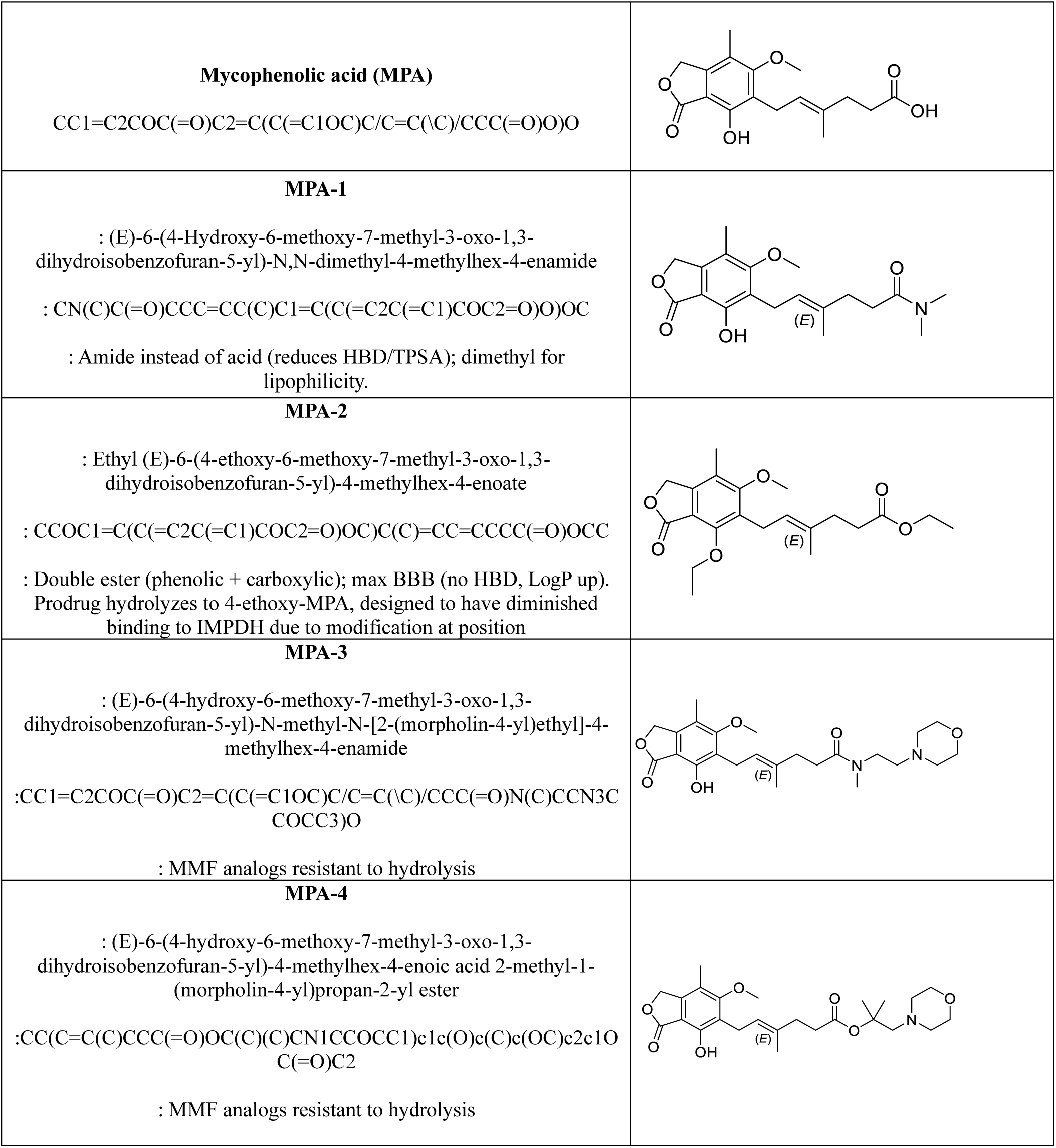
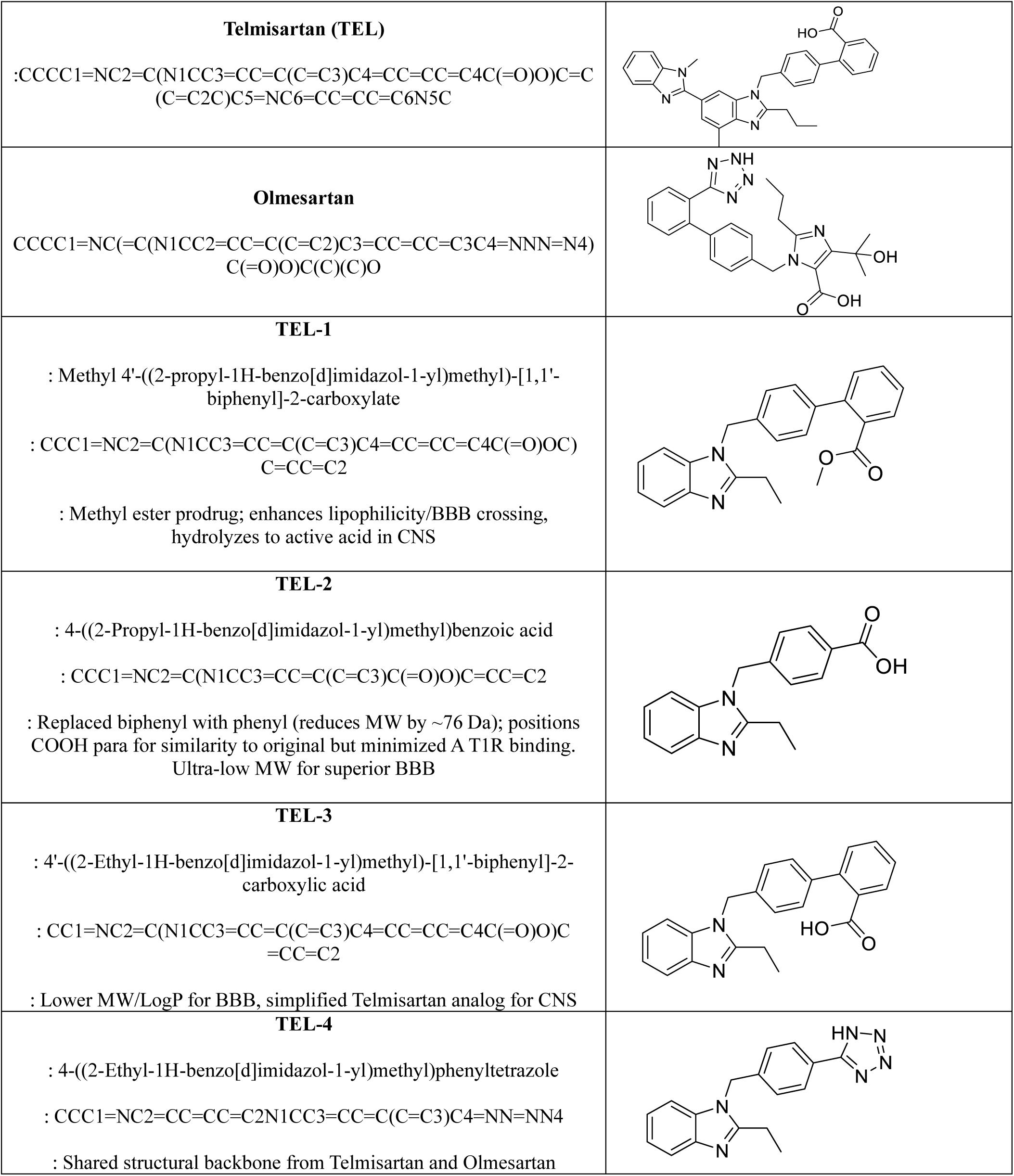
Repurposing FDA-approved drugs and their derivatives to disrupt the CHIP–LMO7–BAG5 inhibitory complex and activate CHIP-dependent tau clearance as a therapeutic strategy for tauopathies.

**Supplementary Figure 1.**
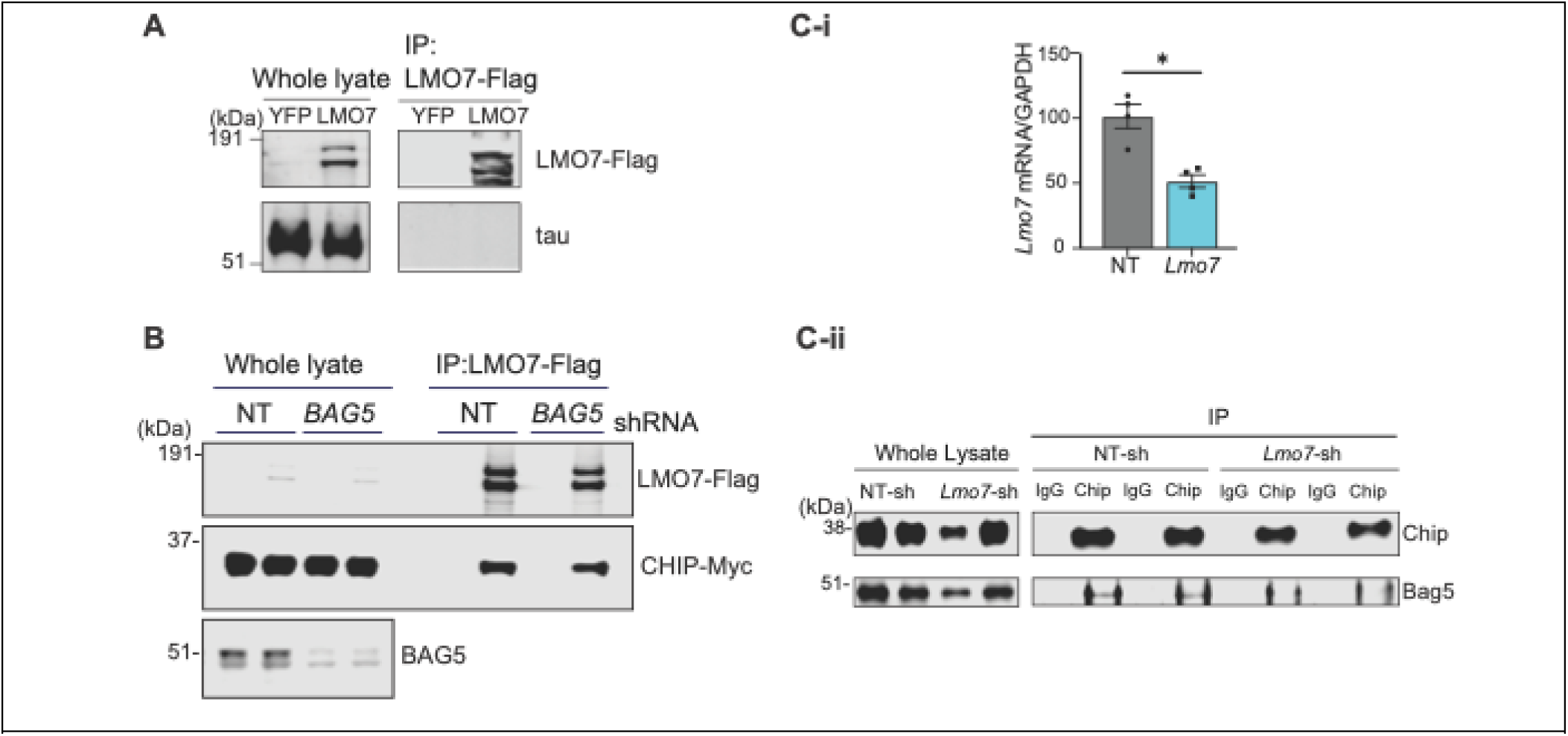
(**A**) IB images showing that LMO7 IP does not pull down tau protein in HEK293T cells overexpressing tau with YFP or LMO7-Flag. (**B**) IB image of co-IP LMO7 by CHIP-IP in HEK293T cells treated with NT shRNA or BAG5 shRNA. **(C-i)** qRT-PCR of *Lmo7* mRNA and **(C-ii)** IB image of co-IP BAG5 by CHIP in mouse brain injected with AAV8 containing either NT shRNA or LMO7 shRNA.

**Supplementary Figure 2.**
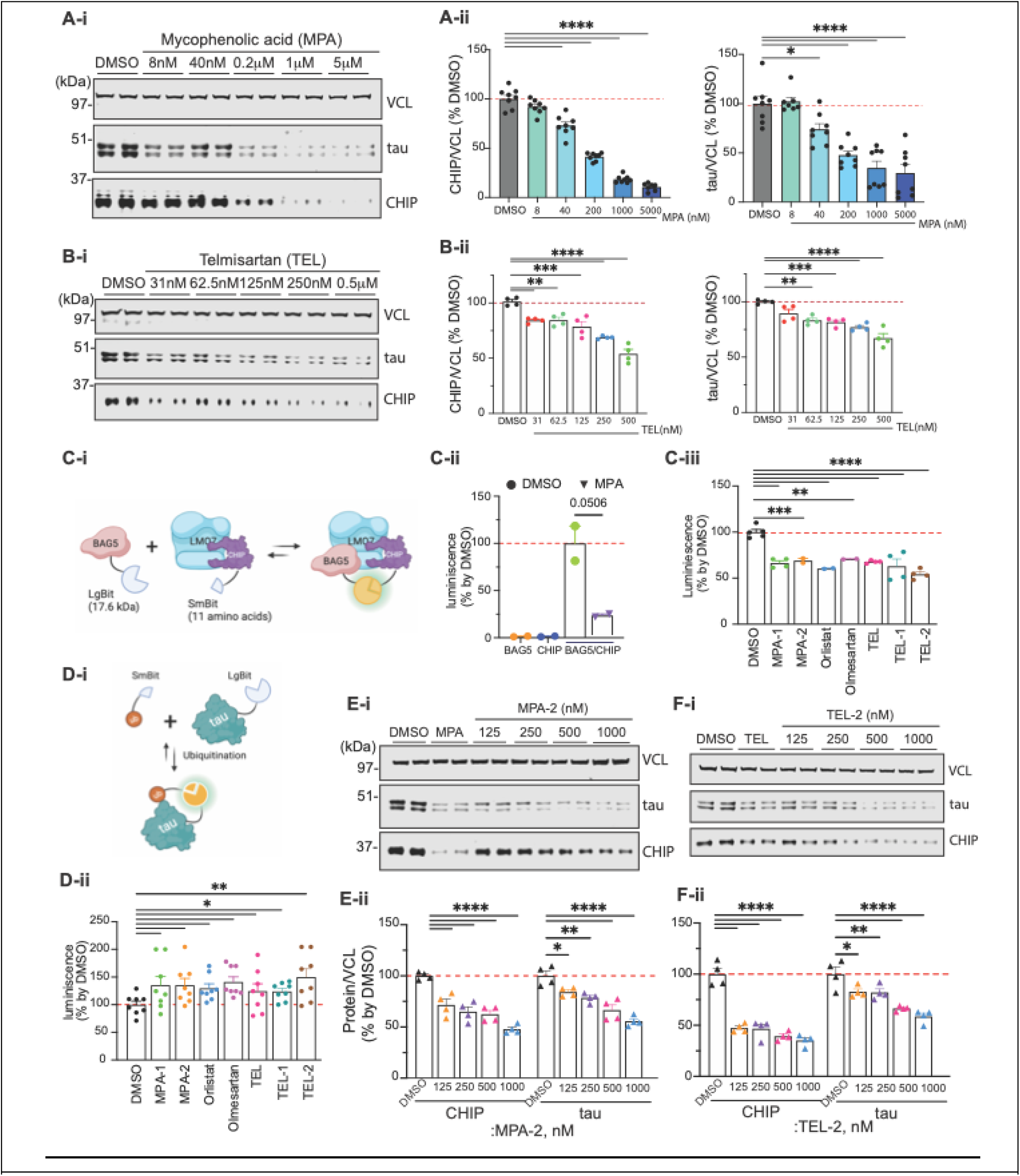
Effects of selected compounds on tau and CHIP levels and BAG5–CHIP interaction. Representative IB image (**A**-**i**) and quantification (**A-ii**) of tau and CHIP levels in HEK293T cells treated with the serial dilution of MPA (0 - 5μM, 48 hours). Representative IB images (B-i) and quantification (B-ii) of tau and CHIP levels in HEK293T cells treated with the serial dilution TEL (0 - 0.5μM, 48 hours). (C-i) NanoBiT assay for measuring the BAG5-CHIP interaction by luminescence signal. (C-ii) Luminescence signal of cells expressing either BAG5-LargeBiT, CHIP-SmallBiT, or both proteins, treated with DMSO or MPA (1μM, 24 hours). (C-iii) Quantification of luminescence signals reflecting BAG5-CHIP interaction in cells treated with DMSO or the indicated compounds (1μM, 24 hours). (D-i) NanoBiT assay to detect mono-ubiquitination of tau proteins indicated by luminescence response by interaction of tau-LargeBiT with ubiquitin-SmallBiT. (D-ii) Quantification graph of tau mono-ubiquitination in live cells treated with the indicated individual compounds (1μM, 24 hours). Representative IB images (E-i and F-i) and quantification graph (E-ii and F-ii) showing the level of tau and CHIP protein in HEK293T cells treated with either MPA-2 (E) and TEL-2 (F) (0 to1μM). Data shown as mean ± SEM (*p≤0.05, **p≤0.01, ***p≤0.001, ****p≤0.0001).

**Supplementary Figure 3.**
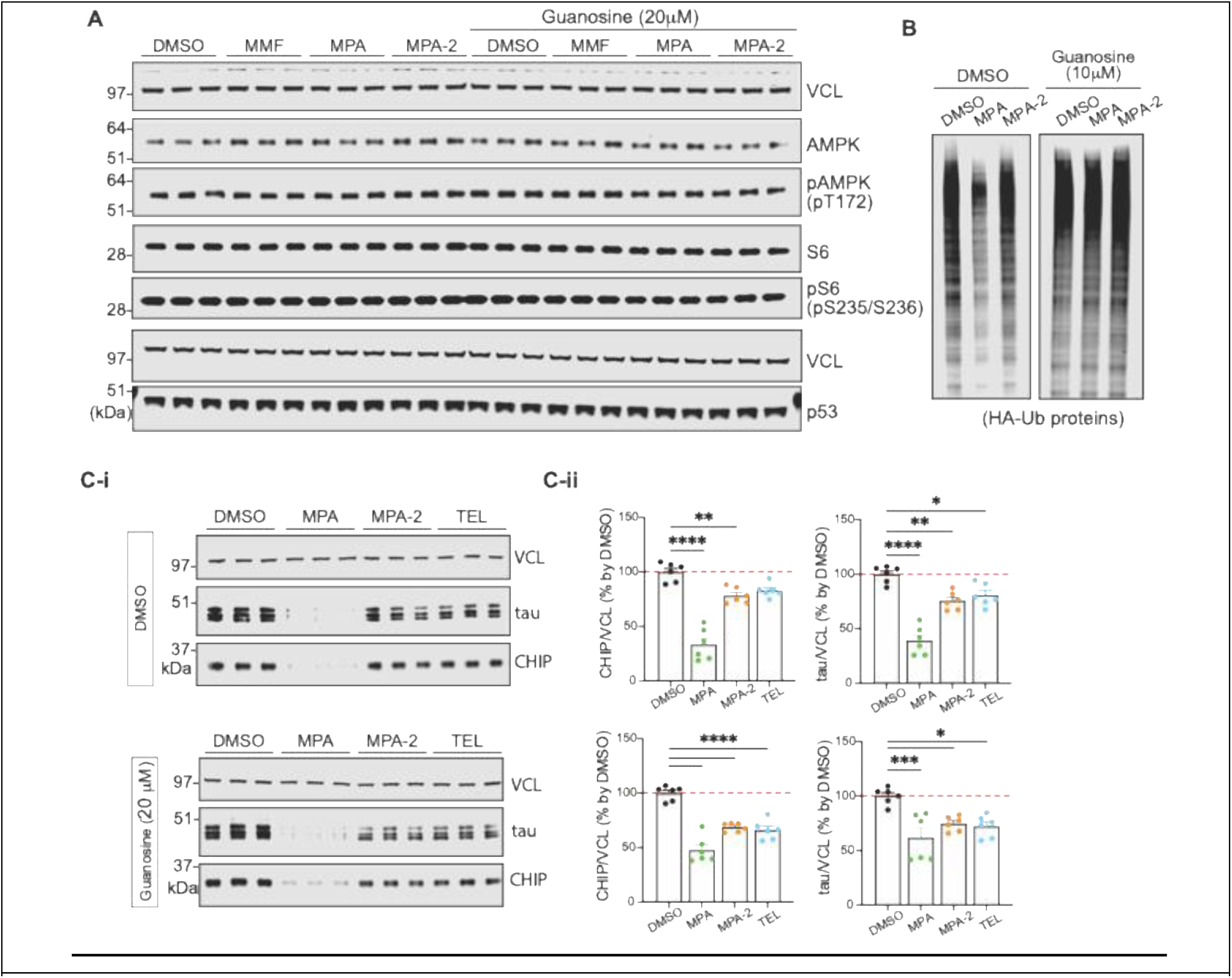
Effects of candidate compounds on cellular stress pathways and tau/CHIP regulation. **(A)** IB image analysis of canonical energy stress, cell growth, and nucleotide stress/apoptosis sensor proteins in HEK293T cells treated with the indicated compounds (1μM). Phospho-AMP-activated protein kinase (p-AMPK) and phosphor-S6 ribosomal protein (p-S6) were analyzed after 6 hours of treatment, and p53 levels were analyzed after 24 hours, in the absence or presence of guanosine supplementation. **(B)** IB analysis of HA-tagged ubiquitinated proteins of whole cell lysate of HEK293T transiently transfected with HA-ubiquitin and treatment with the indicated compounds (1μM, for 48 hours), with or without guanosine supplementation to assess protein abundance. Representative IB images **(C-i)** and corresponding quantification **(C-ii)** of tau and CHIP levels in HEK293T cells treated with the indicated compounds in presence of DMSO (upper panel) or 20 μM guanosine (lower panel). Data shown as mean ± SEM (*p≤0.05, **p≤0.01, ***p≤0.001, ****p≤0.0001).

