## Supplemental data for "The CHIP–LMO7–BAG5 complex controls tau clearance and yields repurposed and newly designed therapeutic candidates for Alzheimer’s disease"

| GeneID | GeneSymbol | p Value | Fold Change LMO7/YFP |
|---|---|---|---|
| 4008 | LMO7 | 0.00 | 29203673 |
| 3416 | IDE | 0.00 | 1830276 |
| 4627 | MYH9 | 0.00 | 712315 |
| 7533 | YWHAH | 0.00 | 553490 |
| 10273 | STUB1 | 0.00 | 395892 |
| 4637 | MYL6 | 0.00 | 278105 |
| 103910 | MYL12B | 0.00 | 241968 |
| 10627 | MYL12A | 0.00 | 217771 |
| 4628 | MYH10 | 0.00 | 197592 |
| 81572 | PDRG1 | 0.00 | 138638 |
| 10471 | PFDN6 | 0.00 | 100562 |
| 8725 | URI1 | 0.00 | 89714 |
| 79657 | RPAP3 | 0.00 | 75697 |
| 8409 | UXT | 0.00 | 69024 |
| 9529 | BAG5 | 0.00 | 68437 |
| 116143 | WDR92 | 0.00 | 61496 |
| 5202 | PFDN2 | 0.00 | 46556 |
| 55011 | PIH1D1 | 0.00 | 44808 |
| 79784 | MYH14 | 0.00 | 29261 |
| 29766 | TMOD3 | 0.00 | 25781 |
| 10728 | PTGES3 | 0.00 | 12601 |
| 7111 | TMOD1 | 0.00 | 10635 |
| 6709 | SPTAN1 | 0.00 | 9678 |
| 81 | ACTN4 | 0.00 | 7119 |
| 6711 | SPTBN1 | 0.00 | 6631 |
| 6421 | SFPQ | 0.00 | 4730 |
| 4430 | MYO1B | 0.00 | 2214 |
| 4644 | MYO5A | 0.00 | 1615 |
| 26160 | IFT172 | 0.00 | 1351 |
| 2317 | FLNB | 0.00 | 737 |
| 369 | ARAF | 0.00 | 697 |
| 7266 | DNAJC7 | 0.03 | 300 |
| 3308 | HSPA4 | 0.03 | 164 |

**Supplementary Table 1. Results of IP-MS showing the proteins that co-precipitated with LMO7.** Fold change indicates the abundance of a specific protein in the LMO7-IP sample compared to YFP control samples. Possible interactors are selected using the cut-off of Fold change >100 and significance p-value<0.05. LMO7 (in red) was efficiently pulled down via our IP and identified in our experimental samples but not in our control YFP. Both STUB1 (CHIP) and BAG5, highlighted in yellow under the Gene Symbol column, were identified as possible interactors of LMO7.

| | |
|---|---|
| **Mycophenolic acid (MPA)**<br><br>CC1=C2COC(=O)C2=C(C(=C1OC)C/C=C(\C)/CCC(=O)O)O | 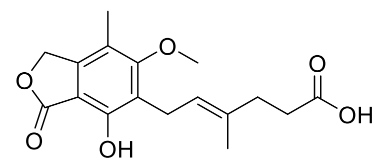 |
| **MPA-1**<br><br>: (E)-6-(4-Hydroxy-6-methoxy-7-methyl-3-oxo-1,3-dihydroisobenzofuran-5-yl)-N,N-dimethyl-4-methylhex-4-enamide<br><br>: CN(C)C(=O)CCC=CC(C)C1=C(C(=C2C(=C1)COC2=O)O)OC<br><br>: Amide instead of acid (reduces HBD/TPSA); dimethyl for lipophilicity. | 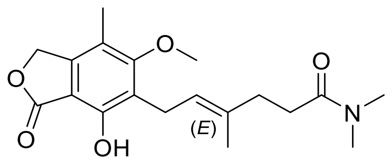 |
| **MPA-2**<br><br>: Ethyl (E)-6-(4-ethoxy-6-methoxy-7-methyl-3-oxo-1,3-dihydroisobenzofuran-5-yl)-4-methylhex-4-enoate<br><br>: CCOC1=C(C(=C2C(=C1)COC2=O)OC)C(C)=CC=CCCC(=O)OCC<br><br>: Double ester (phenolic + carboxylic); max BBB (no HBD, LogP up). Prodrug hydrolyzes to 4-ethoxy-MPA, designed to have diminished binding to IMPDH due to modification at position | 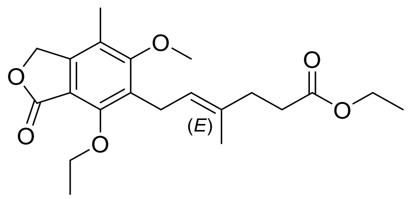 |
| **MPA-3**<br><br>: (E)-6-(4-hydroxy-6-methoxy-7-methyl-3-oxo-1,3-dihydroisobenzofuran-5-yl)-N-methyl-N-[2-(morpholin-4-yl)ethyl]-4-methylhex-4-enamide<br><br>:CC1=C2COC(=O)C2=C(C(=C1OC)C/C=C(\C)/CCC(=O)N(C)CCN3CCOCC3)O<br><br>: MMF analogs resistant to hydrolysis | 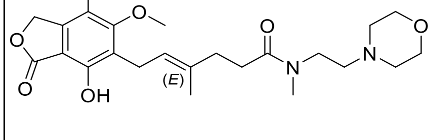 |

| | |
|---|---|
| **MPA-4**<br><br>: (E)-6-(4-hydroxy-6-methoxy-7-methyl-3-oxo-1,3-dihydroisobenzofuran-5-yl)-4-methylhex-4-enoic acid 2-methyl-1-(morpholin-4-yl)propan-2-yl ester<br><br>:CC(C=C(C)CCC(=O)OC(C)(C)CN1CCOCC1)c1c(O)c(C)c(OC)c2c1OC(=O)C2<br><br>: MMF analogs resistant to hydrolysis | 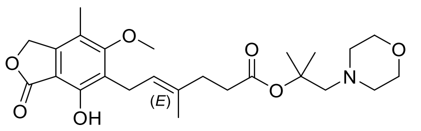 |
| **Telmisartan (TEL)**<br><br>:CCCC1=NC2=C(N1CC3=CC=C(C=C3)C4=CC=CC=C4C(=O)O)C=C(C=C2C)C5=NC6=CC=CC=C6N5C | 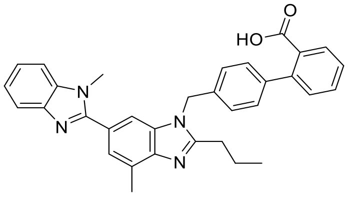 |
| **Olmesartan**<br><br>CCCC1=NC(=C(N1CC2=CC=C(C=C2)C3=CC=CC=C3C4=NNN=N4)C(=O)O)C(C)(C)O | 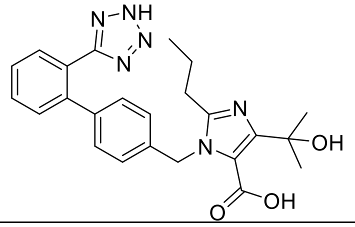 |
| **TEL-1**<br><br>: Methyl 4'-((2-propyl-1H-benzo[d]imidazol-1-yl)methyl)-[1,1'-biphenyl]-2-carboxylate<br><br>: CCC1=NC2=C(N1CC3=CC=C(C=C3)C4=CC=CC=C4C(=O)OC)C=CC=C2<br><br>: Methyl ester prodrug; enhances lipophilicity/BBB crossing, hydrolyzes to active acid in CNS | 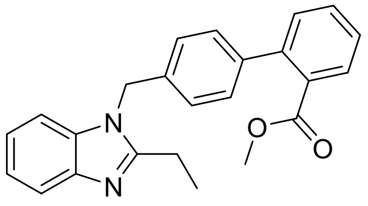 |

| | |
|---|---|
| **TEL-2**<br><br>: 4-((2-Propyl-1H-benzo[d]imidazol-1-yl)methyl)benzoic acid<br><br>: CCC1=NC2=C(N1CC3=CC=C(C=C3)C(=O)O)C=CC=C2<br><br>: Replaced biphenyl with phenyl (reduces MW by ~76 Da); positions COOH para for similarity to original but minimized A T1R binding. Ultra-low MW for superior BBB | 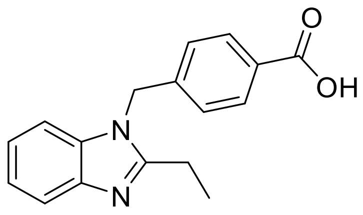 |
| **TEL-3**<br><br>: 4'-((2-Ethyl-1H-benzo[d]imidazol-1-yl)methyl)-[1,1'-biphenyl]-2-carboxylic acid<br><br>: CC1=NC2=C(N1CC3=CC=C(C=C3)C4=CC=CC=C4C(=O)O)C<br>=CC=C2<br><br>: Lower MW/LogP for BBB, simplified Telmisartan analog for CNS | 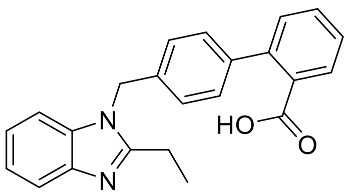 |
| **TEL-4**<br><br>: 4-((2-Ethyl-1H-benzo[d]imidazol-1-yl)methyl)phenyltetrazole<br><br>: CCC1=NC2=CC=CC=C2N1CC3=CC=C(C=C3)C4=NN=NN4<br><br>: Shared structural backbone from Telmisartan and Olmesartan | 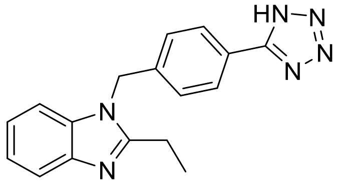 |

**Supplementary Table 2. Repurposing FDA-approved drugs and their derivatives to disrupt the CHIP–LMO7–BAG5 inhibitory complex and activate CHIP-dependent tau clearance as a therapeutic strategy for tauopathies.**

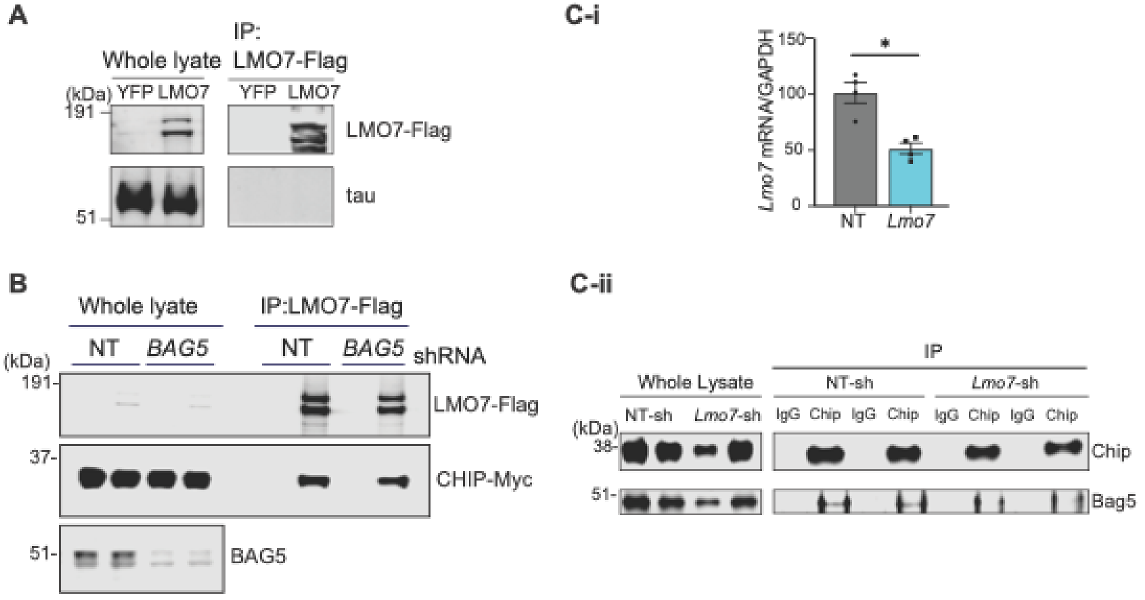


**Supplementary Figure 1.** (**A**) IB images showing that LMO7 IP does not pull down tau protein in HEK293T cells overexpressing tau with YFP or LMO7-Flag. (**B**) IB image of co-IP LMO7 by CHIP-IP in HEK293T cells treated with NT shRNA or BAG5 shRNA. **(C-i)** qRT-PCR of *Lmo7* mRNA and **(C-ii)** IB image of co-IP BAG5 by CHIP in mouse brain injected with AAV8 containing either NT shRNA or LMO7 shRNA.

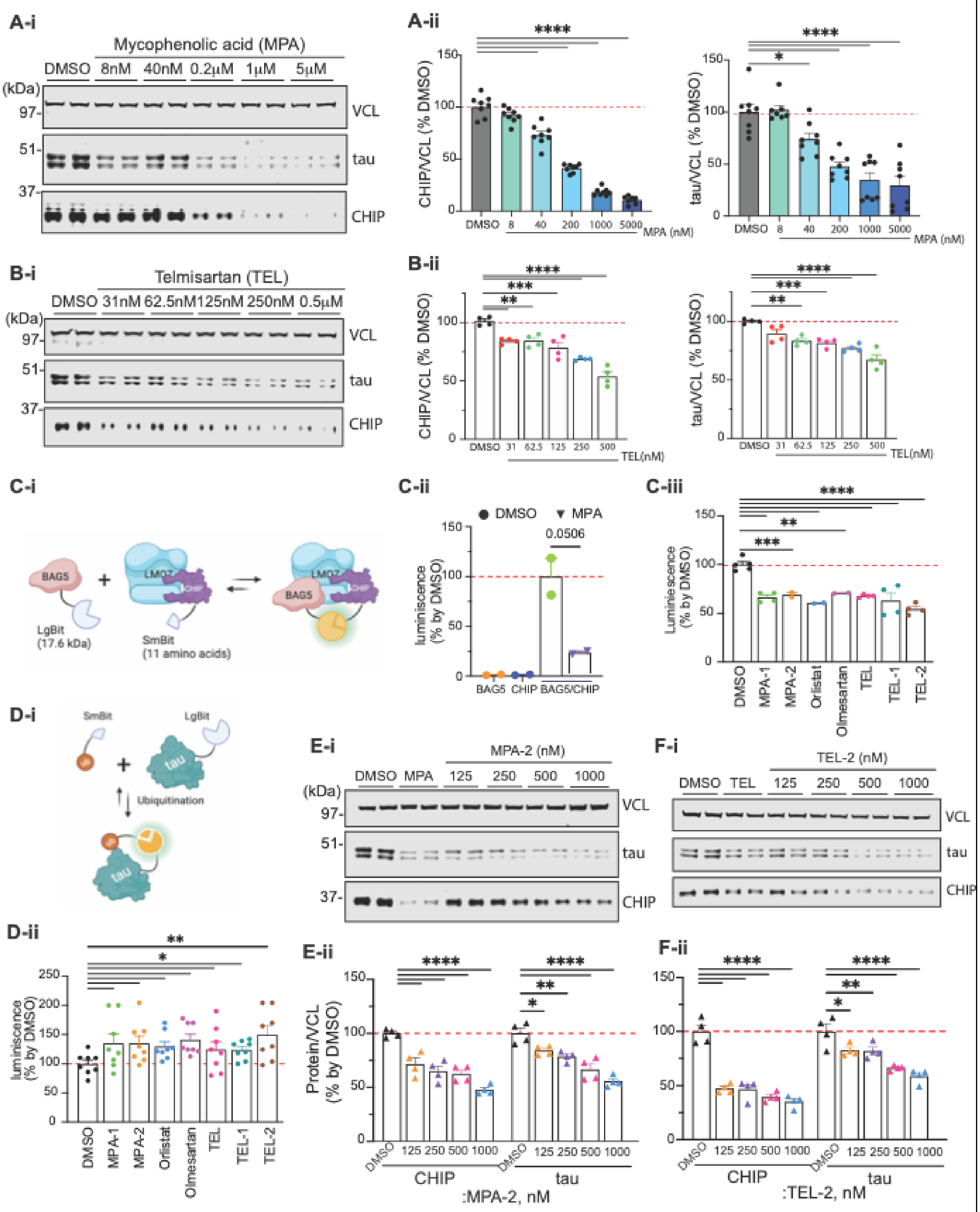


**Supplementary Figure 2. Effects of selected compounds on tau and CHIP levels and BAG5–CHIP interaction.** Representative IB image (**A-i**) and quantification (**A-ii**) of tau and CHIP levels in

HEK293T cells treated with the serial dilution of MPA (0 - 5μM, 48 hours). Representative IB images (**B**-**i**) and quantification (**B-ii**) of tau and CHIP levels in HEK293T cells treated with the serial dilution TEL (0 - 0.5μM, 48 hours). (**C-i**) NanoBiT assay for measuring the BAG5-CHIP interaction by luminescence signal. (**C-ii**) Luminescence signal of cells expressing either BAG5-LargeBiT, CHIP-SmallBiT, or both proteins, treated with DMSO or MPA (1μM, 24 hours). (**C-iii**) Quantification of luminescence signals reflecting BAG5-CHIP interaction in cells treated with DMSO or the indicated compounds (1μM, 24 hours). (**D-i**) NanoBiT assay to detect mono-ubiquitination of tau proteins indicated by luminescence response by interaction of tau-LargeBiT with ubiquitin-SmallBiT. (**D-ii**) Quantification graph of tau mono-ubiquitination in live cells treated with the indicated individual compounds (1μM, 24 hours). Representative IB images (**E-i** and **F-i**) and quantification graph (**E-ii** and **F-ii**) showing the level of tau and CHIP protein in HEK293T cells treated with either MPA-2 (**E**) and TEL-2 (**F**) (0 to1μM). Data shown as mean ± SEM (*p≤0.05, **p≤0.01, ***p≤0.001, ****p≤0.0001).

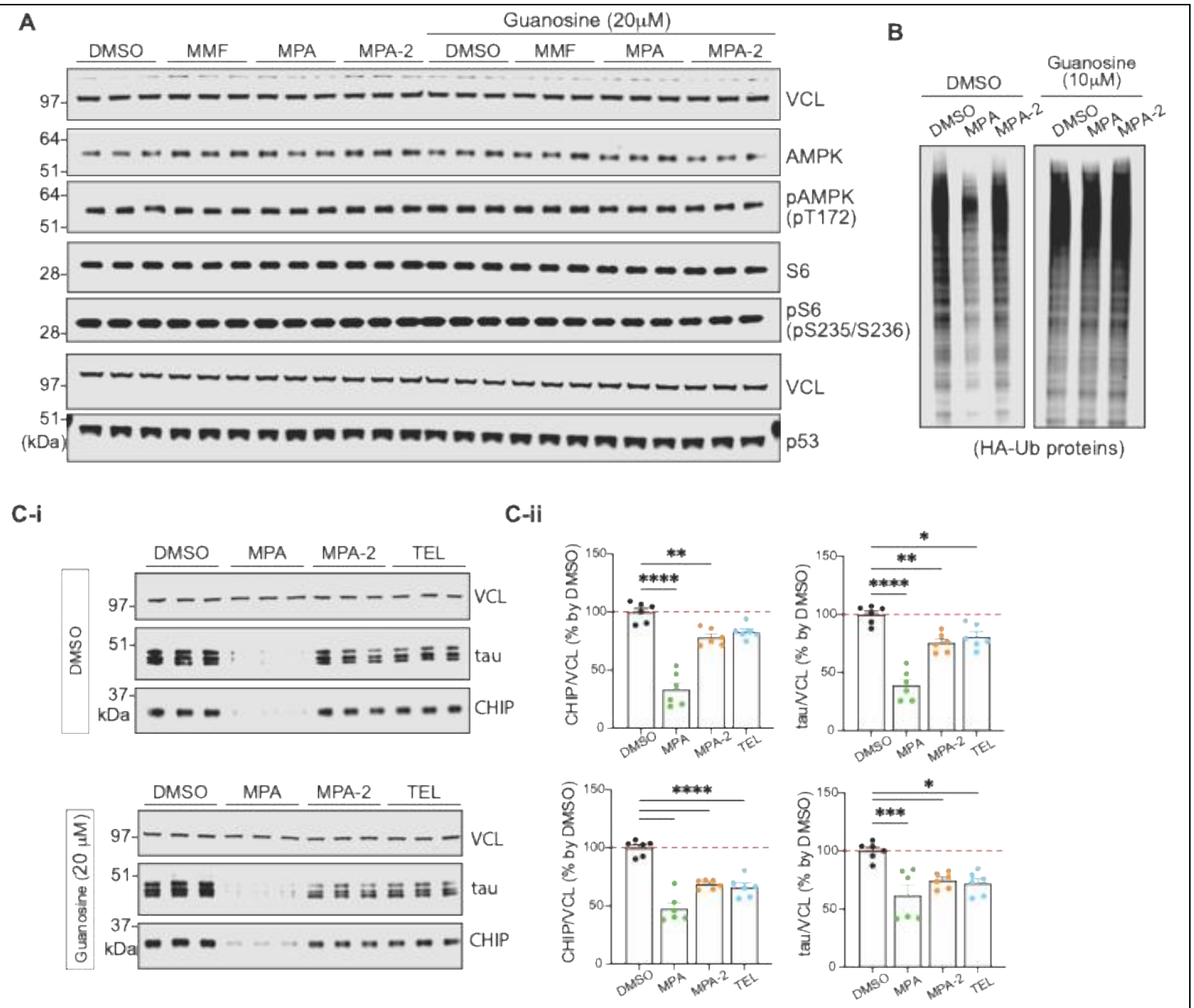


**Supplementary Figure 3. Effects of candidate compounds on cellular stress pathways and tau/CHIP regulation.** **(A)** IB image analysis of canonical energy stress, cell growth, and nucleotide stress/apoptosis sensor proteins in HEK293T cells treated with the indicated compounds (1μM). Phospho-AMP-activated protein kinase (p-AMPK) and phosphor-S6 ribosomal protein (p-S6) were analyzed after 6 hours of treatment, and p53 levels were analyzed after 24 hours, in the absence or presence of guanosine supplementation. **(B)** IB analysis of HA-tagged ubiquitinated proteins of whole cell lysate of HEK293T transiently transfected with HA-ubiquitin and treatment with the indicated compounds (1μM, for 48 hours), with or without guanosine supplementation to assess protein abundance. Representative IB images **(C-i)** and corresponding quantification **(C-ii)** of tau and CHIP levels in HEK293T cells treated with the indicated compounds in presence of DMSO (upper panel) or 20 μM guanosine (lower panel). Data shown as mean ± SEM (*p≤0.05, **p≤0.01, ***p≤0.001, ****p≤0.0001).
